# Cost-effective solutions for high-throughput enzymatic DNA methylation sequencing

**DOI:** 10.1101/2024.09.09.612068

**Authors:** Amy Longtin, Marina M. Watowich, Baptiste Sadoughi, Rachel M. Petersen, Sarah F. Brosnan, Kenneth Buetow, Qiuyin Cai, Michael D. Gurven, Heather M. Highland, Yi-Ting Huang, Hillard Kaplan, Thomas S. Kraft, Yvonne A. L. Lim, Jirong Long, Amanda D. Melin, Jamie Roberson, Kee-Seong Ng, Jonathan Stieglitz, Benjamin C. Trumble, Vivek V. Venkataraman, Ian J. Wallace, Jie Wu, Noah Snyder-Mackler, Angela Jones, Alexander G. Bick, Amanda J. Lea

## Abstract

Characterizing DNA methylation patterns is important for addressing key questions in evolutionary biology, geroscience, and medical genomics. While costs are decreasing, whole-genome DNA methylation profiling remains prohibitively expensive for most population-scale studies, creating a need for cost-effective, reduced representation approaches (i.e., assays that rely on microarrays, enzyme digests, or sequence capture to target a subset of the genome). Most common whole genome and reduced representation techniques rely on bisulfite conversion, which can damage DNA resulting in DNA loss and sequencing biases. Enzymatic methyl sequencing (EM-seq) was recently proposed to overcome these issues, but thorough benchmarking of EM-seq combined with cost-effective, reduced representation strategies has not yet been performed. To do so, we optimized Targeted Methylation Sequencing protocol (TMS)—which profiles ∼4 million CpG sites—for miniaturization, flexibility, and multispecies use at a cost of ∼$80. First, we tested modifications to increase throughput and reduce cost, including increasing multiplexing, decreasing DNA input, and using enzymatic rather than mechanical fragmentation to prepare DNA. Second, we compared our optimized TMS protocol to commonly used techniques, specifically the Infinium MethylationEPIC BeadChip (n=55 paired samples) and whole genome bisulfite sequencing (n=6 paired samples). In both cases, we found strong agreement between technologies (R² = 0.97 and 0.99, respectively). Third, we tested the optimized TMS protocol in three non-human primate species (rhesus macaques, geladas, and capuchins). We captured a high percentage (mean=77.1%) of targeted CpG sites and produced methylation level estimates that agreed with those generated from reduced representation bisulfite sequencing (R² = 0.98). Finally, we applied our protocol to profile age-associated DNA methylation variation in two subsistence-level populations—the Tsimane of lowland Bolivia and the Orang Asli of Peninsular Malaysia—and found age-methylation patterns that were strikingly similar to those reported in high income cohorts, despite known differences in age-health relationships between lifestyle contexts. Altogether, our optimized TMS protocol will enable cost-effective, population-scale studies of genome-wide DNA methylation levels across human and non-human primate species.

## INTRODUCTION

Characterizing and understanding epigenomic variation is important for evolutionary and developmental biology, geroscience, and biomedicine. DNA methylation—the covalent addition of methyl groups to cytosines—is a semi-malleable and environmentally-responsive epigenetic modification integral to gene regulation in many species, including our own (1). Because DNA methylation moderates gene expression throughout the life course, it is critical for processes such as development (2–4), cell programming (5), tissue specificity (6), aging (7–11), and disease progression (12–14). For example, changes in DNA methylation are considered a “hallmark” of the aging process, with most studies reporting age-associated gains in methylation in hypomethylated regions (e.g., promoters and transcribed regions) and age-associated losses in methylation in hypermethylated regions (e.g., heterochromatic regions, Polycomb-repressed regions) (15–17). These age-related patterns are so consistent that DNA methylation patterns have been used to construct molecular clocks that reliably predict chronological age (18,19). Further, because DNA methylation is known to respond to environmental inputs, it has been implicated as a mechanism through which diverse experiences can “get under the skin” to impact long-term physiology and health (e.g., famine (20–24), psychosocial stress (25–29), or infection (30–33)).

To profile genome-wide DNA methylation at scale, most studies rely on reduced representation methods: human studies have largely favored microarrays, while non-human studies have favored reduced representation bisulfite sequencing (RRBS) due to the lack of species-specific microarrays (34). Both methods quantify DNA methylation at a subset (1-5%) of CpGs in the genome, and thus provide a cost-effective strategy relative to genome-wide assays (e.g., whole genome bisulfite sequencing (WGBS)). For example, the Infinium MethylationEPIC v2.0 BeadChip, or EPIC array, covers ∼930K CpG sites including functional elements identified by the ENCODE project (35), DNase hypersensitive sites, and putatively important sites for human disease and development (36,37). In contrast, RRBS fragments DNA using the Msp1 enzyme that cuts DNA at CCGG motifs, which following size selection, enriches for 1-5% of the genome with high CpG content such as CpG islands and gene bodies (34,38). Importantly, both microarrays and RRBS rely on sodium bisulfite, which converts unmethylated cytosines to thymine while leaving methylated cytosines protected from conversion. This chemical reaction requires high pHs and temperatures, which can cause unwanted DNA fragmentation and damage, especially to unmethylated cytosines (39). Ultimately, such damage can create difficulties during library preparation as well as biases in the downstream data (39–41).

Enzymatic methyl sequencing (EM-seq) offers a useful alternative to bisulfite sequencing with several key benefits: EM-seq relies on enzymatic rather than chemical conversion of unmethylated cytosines to thymine, resulting in substantially less DNA damage (40). As a result, whole genome EM-seq has been shown to recover more CpGs sites, have lower duplication rates, have better between-replicate correlations, and require less DNA input than WGBS (40). However, existing EM-seq benchmarked protocols rely on whole genome rather than reduced representation strategies, hindering their adoption especially for population-scale studies. To address this gap, Twist Biosciences recently created a hybrid capture panel that targets ∼4 million CpG sites in the human genome and is compatible with EM-seq. The first generation Twist methylation capture probe set uses ∼550k probes to target functionally relevant CpG sites (e.g., those in enhancers, gene bodies, and near transcription start sites) and to cover 95% of CpG sites included on the widely used EPIC array (42–45). Importantly, off the shelf, this protocol is similar or lower in cost to existing reduced representation approaches. For example, it provides coverage of approximately four times as many CpG sites relative to the EPIC array at one fourth the cost—a ∼16-fold gain in the data-to-price ratio.

Here, we aimed to develop and benchmark an optimized and further cost-reduced version of the targeted methylation sequencing (TMS) approach suitable for population-scale studies, including both human and non-human primate (NHP) studies (Figure 1A). To do so, we built upon the off the shelf TMS protocol (Figure 1B), which recommends 8 plexing of samples per capture reaction and 200 ng of DNA input, and tested four multiplexing strategies (12, 24, 48, and 96 plex, using 200 ng of sample input; Figure 1C). We also tested five DNA input amounts (25, 50, 100, 200, and 400ng, using the 12-plex strategy) and other minor protocol modifications such as varying the annealing temperature during hybrid capture and varying the method used for DNA fragmentation (Figure 1C). Following optimization, we assessed: 1) the robustness of our protocol through a direct comparison with the EPIC array and WGBS; 2) the extension of optimized TMS for use in NHP species; and 3) the utility of our protocol to uncover biological effects of interest (see Table 1 for sample sizes and sample information; Figure 1C). Specifically, we tested for age effects on DNA methylation in two non-industrial populations that exhibit minimal evidence for age-related increases in cardiometabolic diseases—the Tsimane of Bolivia and the Orang Asli of Peninsular Malaysia (46–48). We were interested in the degree to which these groups exhibit epigenomic patterns similar to what has been reported for Western, post-industrial settings, or alternatively, if they exhibit novel patterns that may be associated with their favorable aging trajectories. Overall, we found that we were able to miniaturize and optimize the TMS protocol to ∼$80 per sample, while maintaining data quality and comparability to existing methods, extending to NHP species, and applying the protocol to uncover age effects in two under-represented human populations.

**Figure 1.**
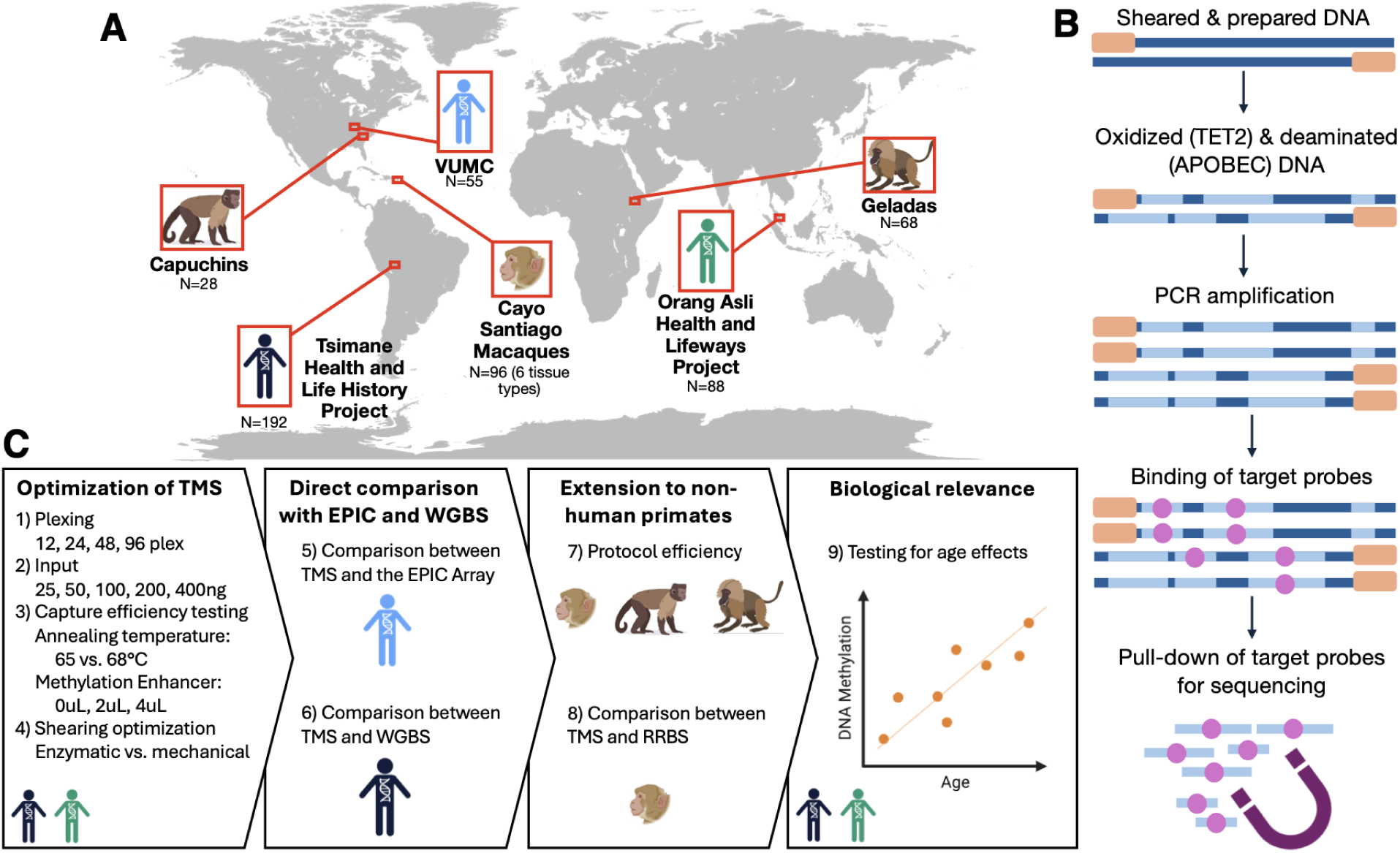
Experimental design and study populations. **(A)** To optimize the TMS protocol, we used samples from three human and three NHP populations: the Tsimane of Bolivia, a Vanderbilt University Medical Center cohort, the Orang Asli of Malaysia, rhesus macaques from Cayo Santiago in Puerto Rico, capuchin monkeys from captive sites throughout the United States, and gelada monkeys from Ethiopia. **(B)** The TMS protocol begins with DNA shearing and adapter ligation. Next, two enzymes, TET2 and APOBEC, are used to oxidize and deaminate the DNA. TET2 recognizes methyl groups attached to cytosines and converts them to Ca/g. APOBEC follows TET2 and converts the unmethylated cytosines to uracils. Following PCR amplification (which converts uracils to thymines), hybrid capture is used to enrich for targeted regions of the genome. Samples are then submitted for high throughput sequencing. **(C)** Overview of experiments and analyses. The samples used for each set of experiments are noted by a population-specific icon.

**Table 1:**
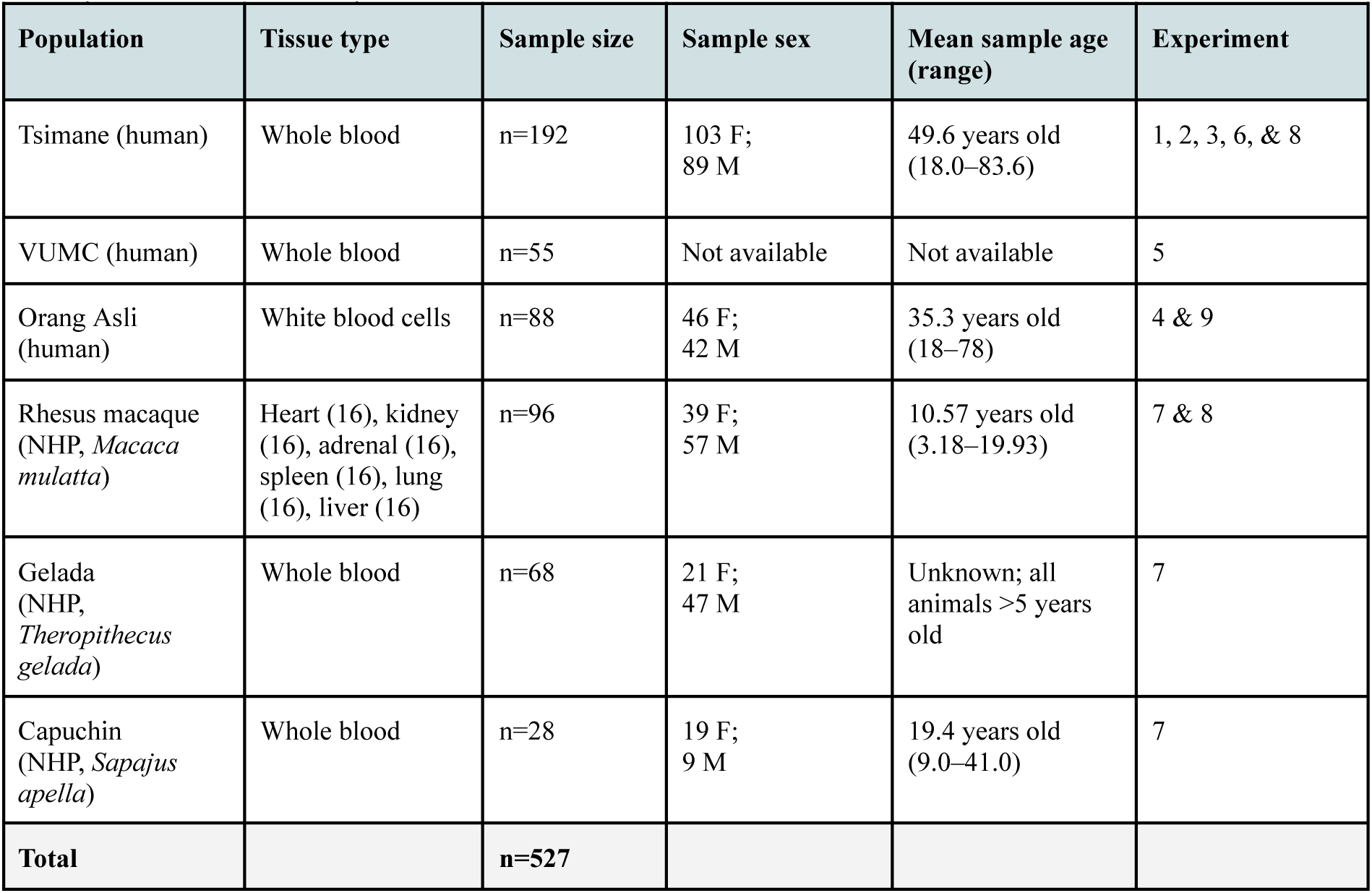
Study populations and sample information for each experiment (names of experiments are as described in Results). NHP = non human primate, F = female, M = male, VUMC = Vanderbilt University Medical Center. See also Table S1 for sample metadata and read depth.

## RESULTS

### Data quality is robust to a range of multiplexing strategies, input amounts, and protocol modifications

#### Experiments 1 & 2: Varying multiplexing strategies and input amounts

Using DNA from Tsimane individuals, we tested four multiplexing strategies (12, 24, 48, and 96 plex, using 200ng of sample input) and five DNA input amounts (25, 50, 100, 200, and 400ng, using the 12-plex strategy). Raw quality control metrics such as percent CHH methylation (a proxy for the rate at which unmethylated cytosines are converted to thymine) and mapping efficiency (percent of reads uniquely mapped to the genome) were high for all samples. Mapping efficiency was consistent across plexing strategies (average mapping efficiency: 12-plex = 71.9%, 24-plex = 72.9%, 48-plex = 72.5%, and 96-plex = 73.5%; ANOVA: F-value = 0.843, p-value = 0.472; Figure 2A) but affected by input amount, with higher DNA input having greater higher mapping efficiency (ANOVA: F-value = 13.57, p-value < 0.001, Figure 2B, Table S2). CHH methylation was consistently well below 1%, indicative of high conversion rate across all plexing and input strategies (range=0.1-0.27%; Figure S1, Table S3 & S4) (49).

**Figure 2.**
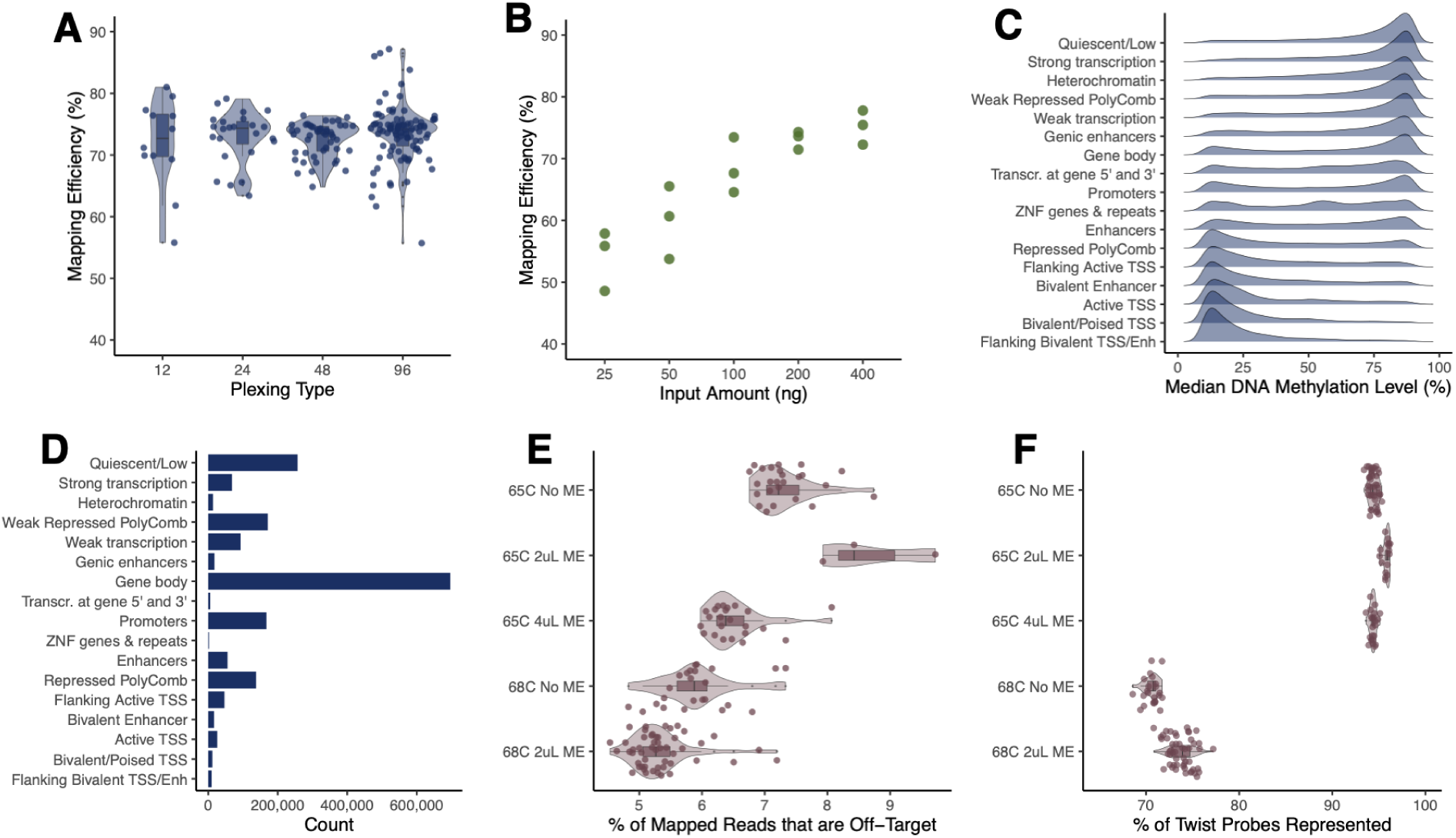
Optimized TMS produces high-quality DNA methylation data across a range of plexing strategies, input amounts, and protocol modifications. **(A)** High (>70%) mean mapping efficiency across plexing strategies. Each point represents a sample within a plexing strategy and the y-axis represents the percent of reads uniquely mapped per sample. **(B)** Mapping efficiency increases as input amount increases. Each point represents a 12-plex pool made with varying DNA input amounts per sample, the y-axis represents the percent of reads uniquely mapped per sample. **(C)** Median DNA methylation levels for reads located within different chromHMM genomic annotations from NIH Roadmap Epigenomics (using data from the 96-plex, 200 ng input from experiment 1). **(D)** The number of CpG sites falling within different chromHMM genomic annotations (using data from the 96-plex, 200 ng input from experiment 1). **(E)** Percent of reads that are not within Twist probes (off-target reads) following protocol modifications to annealing temperature and methylation enhancer (ME) volume. For each set of protocol conditions, the x-axis represents the percent of mapped reads that do not overlap with the Twist probe set. **(F)** Percent of Twist probes that are represented following protocol modifications to adjust the annealing temperature and ME volume. For each set of protocol conditions, the x-axis represents the percentage of Twist probes that were represented by at least 1 read.

After filtering for CpG sites with >5x coverage that were within the Twist probe set (+/- 200bp) and that were covered in the majority of samples in a given experiment, we retained an average of 4,197,008 CpG sites (s.d.=546,767) across plexing experiments and 4,051,941 CpG sites (s.d. = 93,106) across input experiments (Tables S5 & S6). On average, this represented 96.42% and 92.19% coverage of the TMS probe set across the plexing and input experiments, respectively (Table S7 & S8). In addition to consistently recovering the expected set of CpGs, we also observed extremely repeatable methylation levels across the plexing and input experiments (all R^2^>0.99; Table S9 & S10). The CpGs covered by our experiments were distributed across diverse genomic annotations, and the median DNA methylation levels within a given annotation displayed expected patterns (Figure 2C & D) (50). For example, we observed high levels of methylation in quiescent and heterochromatin regions and low levels of methylation in promoters and transcribed regions.

#### Experiments 3 & 4: Optimizing capture efficiency and DNA shearing strategies

In experiments 1 and 2, we used the recommended 65°C annealing temperature during the hybrid capture step—where prepared DNA is bound to the capture probe set to select CpG sites of interest—and the recommended 2uL of methylation enhancer, which increases the efficiency of this reaction. Here, we found that ∼3/4 of all of our mapped reads were “on-target”, meaning that they overlapped with the designed probe set and represented successful hybrid capture (Tables S7 & S8). This suggests that ∼¼ of reads are “off target” and randomly distributed across the genome rather than within our regions of interest (Tables S9 & S10). We therefore performed a third experiment to test two protocol modifications that might decrease the off-target proportion: we increased the annealing temperature (testing 65°C or 68°C) and we varied the amount of methylation enhancer (testing 0uL, 2uL, or 4uL).

In experiment 3, we found that increasing the annealing temperature from 65°C to 68°C resulted in a lower proportion of off-target reads (ANOVA: F-value = 84.2, p-value < 0.0001; Figure 2E, Figure S2, Table S11). Across samples annealed at 65°C, an average of 78.5% of reads were on-target, while this number rose to 84.2% at 68°C. However, this increase in capture efficiency came at a cost to the breadth of CpG sites covered: across samples annealed at 65°C, we observed coverage of on average 92.0% of the probe set, while this number fell to 72.2% for samples annealed at 68°C (Figure 2F; Tables S12 & S13). This suggests that higher annealing temperatures lead to greater but more specific binding during the hybrid capture step, and thus the increased capture efficiency comes at the expense of recovering all the expected CpG sites. In general, we did not find meaningful differences across methylation enhancer amounts and we therefore excluded this reagent from downstream experiments (Figure 2E & F). Given the loss of certain genomic regions at 68°C, downstream experiments focused on a 65°C annealing temperature.

We next performed a fourth experiment focused on protocol optimization, in which we varied the strategies used to fragment genomic DNA prior to EM-seq library preparation: specifically, we tested mechanical shearing via Covaris sonication against enzymatic shearing with the NEBNext UltraShear reagent. Mechanical shearing is the current standard approach but is expensive, requires special equipment, and is labor intensive. Conversely, enzymatic fragmentation is cheaper, does not require special equipment, and is more compatible with automation. For experiments 3 and 4, we used the 96-plex strategy and 200 ng of sample input, since experiments 1 and 2 suggested that data quality does not suffer from higher plexing strategies.

Enzymatic shearing resulted in a similar number of covered sites as was previously observed with mechanical shearing (n=4,591,123 and 4,523,981 filtered CpG sites for the 10 and 20 minute protocols, respectively). Average site-specific methylation levels were also highly concordant between approaches (mechanical versus 10 min enzymatic: R^2^ = 0.9875; mechanical versus 20 min enzymatic: R^2^=0.9876; 10 min versus 20 min enzymatic: R^2^=0.9944; Figures S3 & S4). This was also true when we focused on a subset of paired samples processed using both methods (n=3; mechanical versus 10 min enzymatic: average R^2^ = 0.971; mechanical versus 20 min enzymatic: average R^2^=0.971; 10 min versus 20 min enzymatic: average R^2^=0.987, Table S14). From these experiments, we concluded that enzymatic fragmentation can be substituted into the protocol with no loss to data quality.

We also used these data, which represent our “best” protocol (96-plex, 200ng input, 65°C annealing, no methylation enhancer, enzymatic shearing), to understand a critical aspect of experimental design—how many reads one would need to generate to achieve a given average coverage per CpG site (Figure S5). In general, we observe a 1:2 relationship between the number of mapped, paired end reads (in millions) and average coverage, such that ∼25M mapped paired end reads translates to ∼50x average coverage per CpG site.

### Epigenomic profiles measured with TMS recapitulate those measured with the EPIC array and WGBS

#### Experiment 5: Comparison of TMS to the EPIC array

To ensure that TMS could perform comparably to the most popular current reduced-representation technology (the EPIC array), we generated paired data for 55 samples using both platforms (and following the 96-plexing, 200 ng input TMS protocol from experiment 1). After filtering, we analyzed 682,295 CpG sites common to both technologies, and found high concordance between per-site DNA methylation levels averaged across all individuals in the dataset (R^2^ = 0.97; Figure 3A). This remained true when we subsetted to 235,234 variably methylated sites (i.e., sites with methylation levels >10% and <90%; mean R^2^ = 0.83; Figure 3B). Because methylation patterns are relatively consistent within the human genome, we also confirmed that these correlations were higher for EPIC-TMS data generated from the same sample compared to EPIC-TMS comparisons made between random pairs of samples (mean R^2^ for non-variable sites: 0.92, mean R^2^ for variable sites only: 0.75; Figure S6).

**Figure 3.**
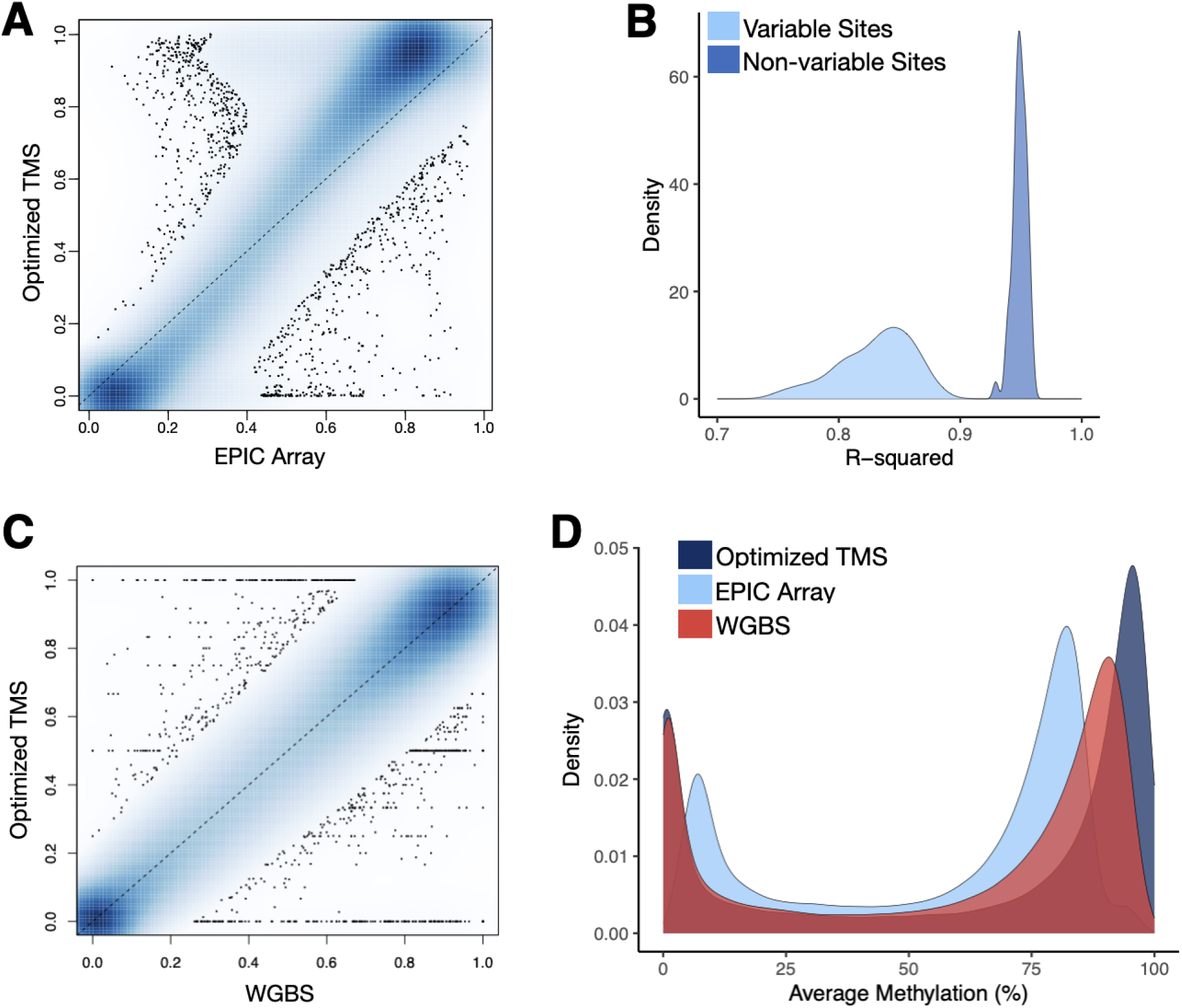
Optimized TMS recapitulates DNA methylation levels measured with the EPIC array and WGBS. **(A)** Correlation in DNA methylation levels for EPIC array versus TMS (R^2^=0.97). Site-level methylation averaged across 6 samples measured using the EPIC array (x-axis) and 96-plex, 200 ng input TMS (y-axis). Each point represents a site measured across both technologies and R^2^ value was generated using linear modeling. Sites filtered to >5X coverage in >75% of samples within each technology. **(B)** Histogram of R^2^ values calculated for each individual sample (i.e., comparing DNA methylation levels measured on both technologies for a given sample). R^2^ values are provided when all CpG sites common to both technologies are included, as well as when only variably methylated CpG sites are included. **(C)** Correlation in DNA methylation levels for WGBS versus TMS (R^2^=0.9871). Site-level methylation averaged across 6 samples measured using WGBS (x-axis) and 96-plex, 200 ng input TMS (y-axis). Each point represents a site measured across both technologies and R^2^ value was generated using linear modeling. Sites filtered to >5X coverage in >75% of samples within each technology. **(D)** Density plot of the average DNA methylation levels detected for matched sites between the three technologies (713,282 sites). Notably, the EPIC array is biased against DNA methylation levels of 100%, as previously observed by Shu et al. (2021) (51) and explained by the equation used to calculate beta values.

Of note, these analyses reconfirmed a known bias in the EPIC array data (51), which does not allow for methylation levels of 100% (such that the correlation between average TMS- and EPIC-measured DNA methylation levels is slightly off the x=y line; Figure 3A). This is because EPIC-derived DNA methylation levels are represented as beta values, calculated as the ratio of the intensity of the methylated bead type to the total locus intensity plus an offset value. Due to the addition of the offset value, beta values of 1 are mathematically impossible.

#### Experiment 6: Comparison of TMS to WGBS

For further validation, we also generated WGBS data for 6 samples included in experiment 3 (96-plexing, 200 ng input, 65°C annealing temperature, no ME, mechanical fragmentation). After filtering and merging with the TMS data, we retained 3,078,771 CpG sites covered by both the TMS and WGBS approaches. For these sites, the average methylation levels observed across technologies was highly correlated (R^2^: 0.9871; Figure 3C). We also found that the genome-wide distribution of DNA methylation levels derived from WGBS was more similar to TMS than to the EPIC array, specifically in that it included many sites with average methylation levels of 100% or close to 100%, as expected (Figure 3D, Figure S7).

### TMS can be effectively applied to non-human primate species

#### Experiment 7: Applying TMS to capuchin, rhesus macaque, and gelada samples

To enable epigenomic analyses in our close primate relatives, we also tested whether TMS (96-plex, 200ng input protocol from experiment 1) could be effectively applied to three NHP species: capuchins (*Sapajus apella*; n=28 samples from blood), rhesus macaques (*Macaca mulatta*; n=96 samples from 6 tissues (see Figure S8 and Table S15), and geladas (*Theropithecus gelada*; n=68 samples from blood). While the probe set is designed from the human genome, NHP species share high levels of sequence homology with humans, especially in coding regions and regions near genes (52), leading us to hypothesize that a majority of CpG sites would be recovered. We mapped the Twist probe sequences to each of the NHP genomes to confirm this intuition, and from this analysis expected to capture 3.0-4.8 million CpG sites across the three species (Figure 4B). Importantly, for the rhesus macaque samples, we also generated paired RRBS data and compared our TMS results to a technology that does not rely on hybrid capture.

**Figure 4.**
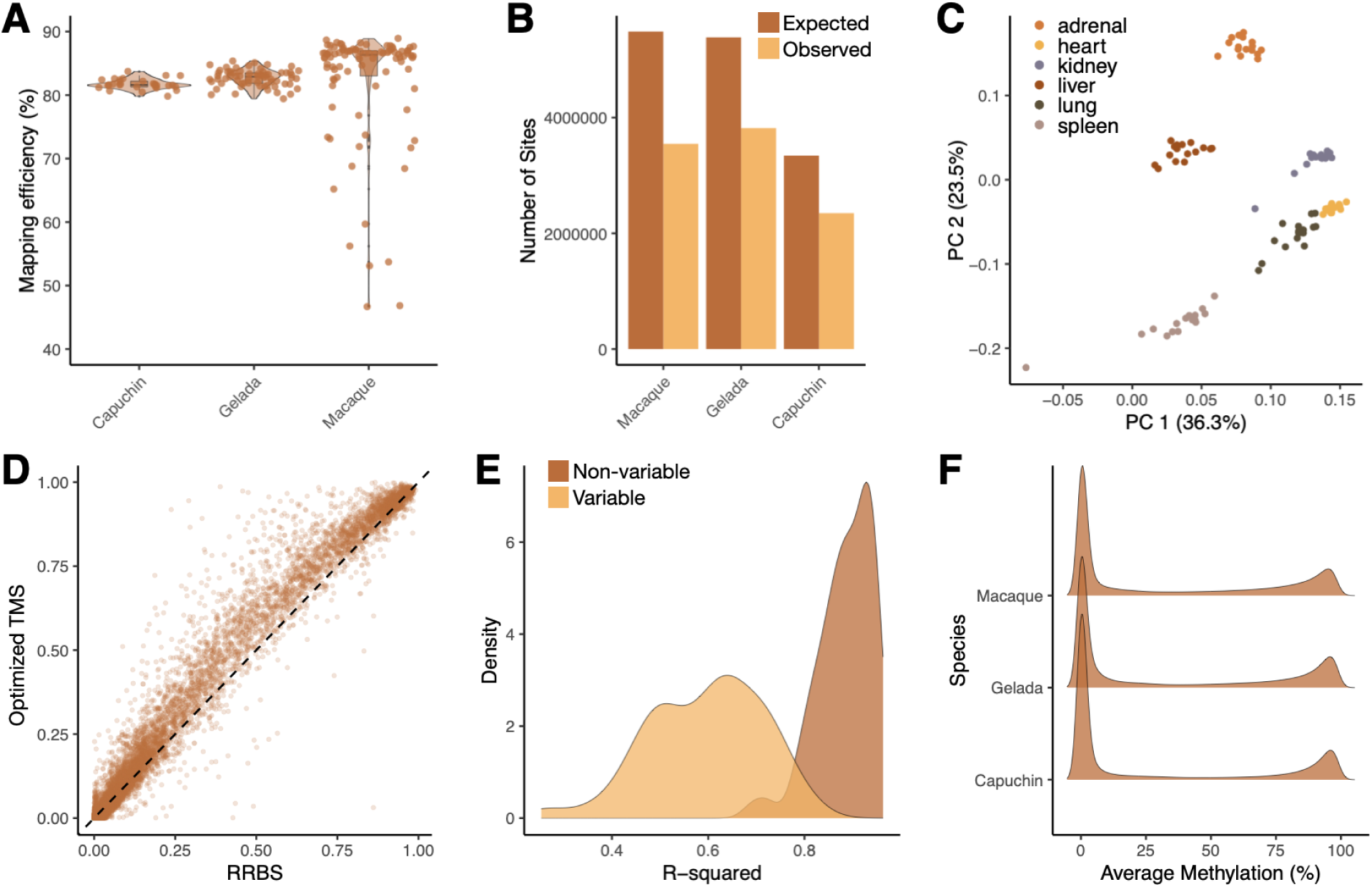
Optimized TMS performs well in non-human primate species and when compared to RRBS. **(A)** Optimized TMS in NHPs results in high mapping efficiencies despite the use of human-specific probes. Here, each of the species are mapped to their own genome. TMS data from experiment 1 (96 plex, 200ng input) are included for comparison. We hypothesize that low mapping efficiency in certain rhesus macaque samples is due to variations in types of tissues processed. **(B)** Number of expected and observed CpG sites covered in each NHP genome. Expected sites were derived from mapping the Twist probes to each NHP genome, while observed sites represent those detected with an average coverage >5X in >75% of samples. **(C)** Principal components analysis of TMS-derived DNA methylation levels from rhesus macaque samples spanning six distinct tissues. **(D)** Similar levels of DNA methylation are detected on a site-by-site basis using RRBS (x-axis) and optimized TMS (y-axis) (R^2^ = 0.97). **(E)** Density plot of LM R^2^ when comparing data generated via optimized TMS and RRBS for the same rhesus macaque sample. R^2^ values are provided when all CpG sites common to both technologies are included, as well as when only variably methylated CpG sites are included. **(F)** Density curves of the average methylation calculated at a given site across samples for each of the NHP species. Curves show expected bimodal distribution in which much of the CpG sites are either hypomethylated or hypermethylated.

When examining initial quality control metrics, we found that all three NHP species had high mapping efficiencies (average = 81.96% for capuchins, 82.62% for geladas, and 81.35% for macaques; Figure 4A). Further, the average CHH methylation levels were all extremely low (<1%), again suggesting high conversion rates (Figure S9). Following filtering, we recovered ∼½ to ¾ of expected CpG sites in the NHP datasets (3,343,133 in capuchin, 5,387,280 in gelada, and 5,486,073 in rhesus macaque). The number of sites recovered scales generally with divergence time (capuchins share a common ancestor with humans 35-45 million years ago, geladas and rhesus macaques share a common ancestor with humans 23-28 million years ago) (53). In all species, we were able to reliably measure more sites than would be typical of RRBS (see below), and we note that some of the between-species variation in performance could be explained by heterogenous read depth (Table S16) as well as reference assembly quality.

When examining average DNA methylation levels across species, we found that, as expected, all exhibited bimodal genome-wide profiles similar to humans (Figure 4F). Further, because the rhesus macaque samples were derived from 6 different tissue types (Figure S8, Table S15), we also confirmed that samples displayed expected tissue-specific epigenetic patterns. Specifically, we demonstrated that a Principal Components Analysis (PCA) was able to reliably separate samples by tissue type (Figure 4C), as has been observed in previous studies using both bisulfite sequencing and the EPIC array (54–56).

#### Experiment 8: Comparison of TMS to RRBS

Studies of NHP species have historically relied on RRBS because of the species-specificity of microarray technologies and the cost barrier of WGBS (57–59). To test how our optimized TMS protocol compares to RRBS, we generated paired data for all 96 rhesus macaque samples. After filtering both datasets to 721,766 common CpG sites, we found a high concordance of the average DNA methylation levels estimated by both technologies (R^2^=0.97; Figure 4D, Figure S10). This remained true when we subsetted specifically to 92,692 variably methylated CpG sites (i.e., sites with average DNA methylation levels >0.1 and <0.9; R^2^=0.5945; Figure 4E).

### TMS reveals consistent age-associated DNA methylation patterns in diverse populations

We next sought to test whether TMS could reliably detect biological signatures of aging and—because we generated data from several human populations—provide novel insights into how diverse environments and ecologies affect human molecular aging. Much of the human data we generated in optimizing TMS came from two Indigenous, non-industrial, subsistence-level populations—the Tsimane of Bolivia, who practice small-scale horticulture and foraging, and the Orang Asli of Peninsular Malaysia, who are comprised of multiple ethnolinguistic subgroups practicing mixed subsistence strategies (60–63). Both groups have participated in long-term, integrated studies of anthropology and health (47,48), which have revealed minimal evidence for age-associated increases in non-communicable diseases such as cardiovascular disease, obesity, and hypertension that are common in high income, post-industrial contexts (64,65). Consequently, we were interested in understanding the degree to which age-associated DNA methylation patterns in these groups broadly recapitulate what has been reported in cohorts from high-income countries. More generally, very little population-scale work has characterized epigenomic aging in populations outside of high-income countries (but see (29,66,67)).

When we tested for age effects on DNA methylation levels in each population (focusing on variably methylated CpG sites and using beta-binomial models (68)), we found that 22% of tested CpGs sites (n=229,727, FDR<5%) were significantly associated with age in the Orang Asli (n=88, age range = 18-78). Of the significant age-associated sites, 40% gained methylation with age and 60% lost methylation with age, recapitulating global patterns observed in previous studies (15,69). In the Tsimane (n=94, age range = 18-75), 0.21% of tested CpG sites (n=1,979) were significantly associated with age, with the lower number of age-associated sites potentially driven by the more restricted age distribution in this sample. Specifically, the Tsimane dataset included 83% of samples >40 years old compared to the Orang Asli dataset, which included 31% of samples >40 years. In the Tsimane, we again observed a global bias toward hypomethylation with age (62% of significant age-associated sites).

Next, we compared the age effect estimated for each CpG site in the Tsimane and Orang Asli with age effects estimated from a modern, post-industrial Swedish cohort previously published in (70) (n=421, age range = 14-94, data generated on the Human Methylation450 BeadChip). Across the 64,084 variably methylated CpG sites measured in all datasets, estimates of the age effect were similar (Pearson correlation: Tsimane v. Orang Asli: R = 0.24, p < 10^-16^; Orang Asli v. Swedish: R = 0.59, p < 10^-16^; Tsimane v. Swedish: R = 0.35, p = 0). Among CpG sites that were significantly associated with age (FDR < 10%), we observed even higher correlations in age effects (Tsimane v. Orang Asli: n=259, R = 0.72, p = 1.07 x 10^-42^; Orang Asli v. Swedish: n=20642, R = 0.76, p = 0; Tsimane v. Swedish: n=424, R = 0.66, p = 3.30 x 10^-54^; Figure S11). Interestingly, the overlap of significant age-associated CpGs sites was more consistent between the two subsistence-level populations than between either subsistence-level population and the Swedish cohort (Fisher’s exact test: Tsimane v. Orang Asli: OR = 2.22, CI = 1.8 - 2.7, p = 7.81 x 10^-18^; Orang Asli v. Swedish: OR = 2.18, CI = 2.0 - 2.3, p = 2.71 x 10^-205^; Tsimane v. Swedish: OR = 1.42, CI = 1.06 - 1.94, p = 0.02).

As methylation changes during aging are known to differ by genomic context, we investigated the extent to which age-associated CpG sites were more likely to show gains (hypermethylation) versus losses (hypomethylation) with age within different functional annotations (Figure 5) (59,70). Despite their diverse lifestyles, epidemiological patterns, and genetic backgrounds, we found strikingly similar age patterns between the three populations. For example, all three populations exhibited age-related hypomethylation in quiescent regions and regions weakly repressed by Polycomb complexes, as well as age-related hypermethylation in promoters, bivalent TSS regions, and enhancers (Figure 5; Table S17). However, there were also some functional annotations that showed patterns specific to subsistence-level lifestyles (e.g., enrichment of hypermethylation with age in gene bodies) or to certain populations (e.g., depletion of hypermethylation with age in heterochromatin in Tsimane), suggesting that while some regions of the genome may exhibit “universal” epigenomic changes with age, others may be modified by environmental or lifestyle factors.

**Figure 5:**
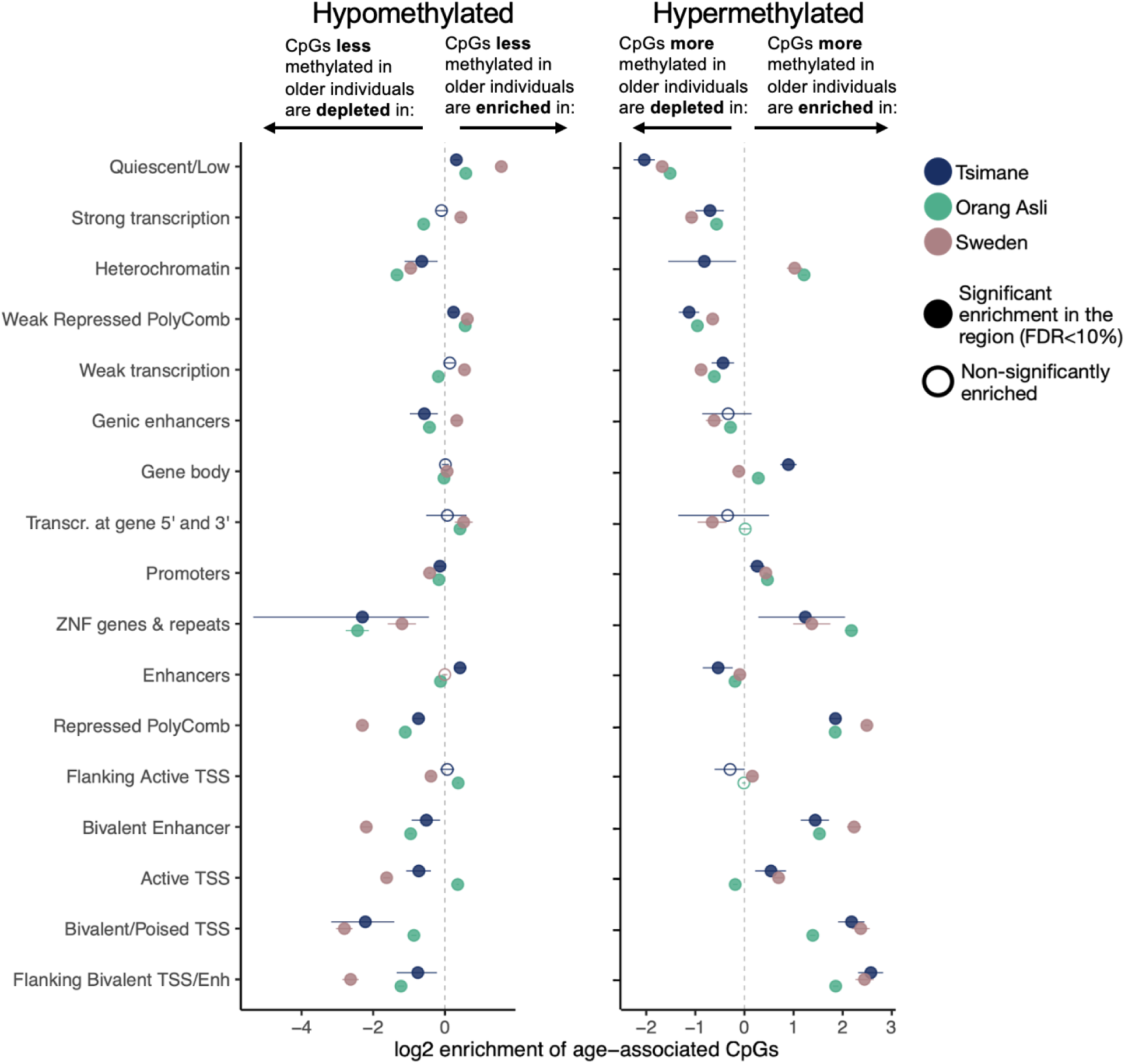
TMS reveals that age-associated DNA methylation patterns are broadly consistent across small scale, subsistence-level populations and post-industrial populations. After quantifying the CpG sites that were differentially methylated with age, we tested the extent to which CpG sites that were hypomethylated with age (i.e., sites with lower methylation levels in older individuals) and hypermethylated with age (i.e., sites with higher methylation levels in older individuals) were more or less likely to fall in different genomic regions. The y-axis is ordered by median methylation levels (as in Figure 2C-D), such that regions higher on the y-axis values have higher average methylation levels regardless of age. We performed enrichment tests for the three populations in which we quantified age effects, the Tsimane (experiment 1 - 96-plex condition, 200ng DNA input, mechanical fragmentation method), the Orang Asli (experiment 4 - 96-plex condition, 200ng DNA input, enzymatic fragmentation method), and a cohort from a high-income country (Sweden) using summary statistics from Johansson et al. (2013).

## DISCUSSION

We developed and thoroughly benchmarked a multiplexed, cost-effective version of the TMS protocol and applied it to diverse human populations as well as multiple NHP species. We recommend an optimal protocol for future work (96-plex, 200ng input, 65°C annealing, no methylation enhancer, enzymatic shearing), but found that data quality remained high across plexing strategies, input amounts, and protocol modifications. Importantly, the 96-plex version of the TMS protocol—including sequencing to achieve ∼50x coverage per CpG site on the Illumina NovaSeq X—can currently be performed for ∼$80/sample (with roughly half being reagents and labor, and the other half being sequencing on the NovaSeq X platform). Relative to the commonly used EPIC array for human studies, which costs ∼$400 per sample and profiles ∼¼ the number of sites covered by TMS, this represents massive savings enabling larger-scale, population-based studies.

We found high concordance between TMS-derived DNA methylation levels and those derived from other commonly used methods—namely the EPIC array, WGBS, and RRBS. WGBS is the gold standard for comprehensive DNA methylation measurement, but is prohibitively expensive for most studies. RRBS has filled in as a more cost-effective alternative, but due to the stochastic nature of the Msp1 digest followed by size selection, not all CpG sites are reliably covered across individuals and missing data can impede downstream analyses (Figure S12). Both WGBS and RRBS also require somewhat specialized laboratory and bioinformatics expertise to execute. As a result of these challenges, microarray-based methods (such as the EPIC array) have become the most commonly used approach in human genomics. Consequently, many popular bioinformatics pipelines and specialized algorithms for DNA methylation data (e.g., epigenetic clocks or cell type deconvolution (72,73)) are currently keyed to microarrays. While DNA methylation levels derived from TMS are strongly correlated with the EPIC array, it is important to keep in mind that: 1) a small subset of sites are not covered by both technologies, and 2) because microarrays output beta values (equivalent to methylated signal/(total signal + an offset)), the relationship between TMS- and EPIC-derived values can not be exactly 1:1 (as noted previously (51)). It is of course possible to recalibrate or rederive popular algorithms such as epigenetic clocks, but we caution that care will be needed when applying microarray-based algorithms to TMS data (as would be true for WGBS or RRBS as well).

The study of DNA methylation in NHP species is deeply important to our understanding of gene regulatory evolution (74–76), comparative aging (58,59,77,78), and environmental impacts on phenotype (59,79). For example, both captive and field-based NHP studies have strongly contributed to our understanding of how social and ecological inputs “get under the skin” to influence fitness-related traits through changes in DNA methylation (80,81). Because microarrays are generally species-specific (but see (82,83)), NHP studies have struggled to identify easy to use, reduced representation approaches for large-scale studies, with most previous work relying on RRBS (59,78,79,81). Although RRBS is easily adapted for non-human species, TMS can work with smaller amounts of input DNA than bisulfite-based protocols (40), which can be critical for studies of wild or endangered species. While TMS uses capture probes designed from the human genome, NHPs share high levels of sequence similarity, which we show is sufficient to reliably capture 2-3 million CpG. Even though not all ∼4 million CpG sites are captured, this still represents a more consistent and cost-effective approach relative to the alternatives. Notably, we find that TMS is effective in both catarrhine (monkeys of Africa and Asia) and platyrrhine (monkeys of Central and South America) species, suggesting it may be effective in other members of these clades.

We applied our optimized TMS protocol to profile DNA methylation levels in two subsistence-level, small-scale, Indigenous populations—the Tsimane of Bolivia and the Orang Asli of Peninsular Malaysia. Both populations have partnered with anthropologists and biologists in long-term studies of behavior, health, and genomics (via The Tsimane Health and Life History Project and The Orang Asli Health and Lifeways Project, respectively (47,48)). These studies have revealed minimal evidence for age-associated increases in cardiometabolic disease in subsistence-level contexts (see also (64,84,85)). Conditions such as cardiovascular disease, type 2 diabetes, and hypertension are widely regarded as being inevitable “diseases of aging” in Western societies (86–88). Further, analyses across a gradient of market-integration and acculturation with the Orang Asli have suggested that lifestyle can directly modify age-related health patterns (89). We therefore wondered to what degree the well-established “hallmarks” of epigenomic aging, derived almost entirely from studies of high-income cohorts, were reflected in these populations (16,90). Our results reveal more similarities than differences in how the epigenomes of the Tsimane, Orang Asli, and high-income individuals change with age. This may reflect recent realizations that much of the epigenome is not functionally important in certain developmental stages and/or cell types, and is thus not actively maintained with age. Instead, DNA methylation patterns throughout much of the genome are thought to decay via a stochastic but predictable process, whereby consistently hypermethylated regions lose methyl marks with age and consistently hypomethylated regions gain methyl marks with age (71,91). However, we find that more functionally important regions of the genome such as enhancers and actively transcribed regions display heterogeneity both in the direction of age effects and how this manifests across populations, leaving clear scope for future studies of environmental and lifestyle effects on epigenomic aging patterns (as well as extensions such as studies of epigenomic age acceleration (66,92,93)). Together, our optimized TMS protocol has the potential to add value and enable larger-scale studies in the many fields that query DNA methylation patterns, such as genetic medicine, developmental biology, evolutionary biology, anthropology, public health, geroscience, and more.

## METHODS

### Study populations, sample collection, and DNA extraction

#### Tsimane of Bolivia

The Tsimane are an Indigenous horticulturalists population spread across >90 villages in the Bolivian lowlands and totaling approximately 17,000 people (47). We extracted DNA from 192 venous whole blood (WB) samples collected between the years of 2010–2021 as part of the Tsimane Health and Life History Project (THLHP) and stored long-term at -80C. The THLHP has continuously collected demographic, behavioral, environmental, and health data along with the provision of medical services for over two decades (94) (University of New Mexico IRB: #07-157; University of California, Santa Barbara IRB: #3-21-0652). The sample set for this project included 103 females and 89 males, with a mean age of 54.3 years old (range 18.0–83.6 years old) (see Table 1). Genomic DNA was extracted using the Zymo *Quick*-DNA 96 kit (Zymo Research #D3012) following the manufacturer’s instructions.

#### Orang Asli of Peninsular Malaysia

The Orang Asli consist of ∼19 ethnolinguistic groups and a total population of ∼210,000 people (48). They traditionally subsist on a mixture of hunting, gathering, fishing, small-scale farming, and trade of forest products (60,61). We extracted DNA from 88 white blood cell (WBC) samples that were collected in 2023 as part of the Orang Asli Health and Lifeways Project (Vanderbilt University IRB #212175). These samples were derived from venous blood draws followed with washing with QIAGEN PureGene red blood cell lysis. Samples were stored in liquid nitrogen upon collection, and at -80C for longer term storage. The Orang Asli sample included 46 females and 42 males, with a mean age of 35.3 years old (range 18–78 years old; Table 1). Genomic DNA was extracted using the Zymo *Quick*-DNA/RNA MagBead kit (Zymo Research #R2131) following the manufacturer’s instructions.

#### Vanderbilt University Medical Center cohort

We were granted access to de-identified EPIC array data (Infinium MethylationEPIC v2.0 Kit) and TMS data from 55 paired human whole blood samples. These samples were sourced from a healthy cohort recruited through the Vanderbilt University Medical Center (VUMC) in Nashville, TN USA. Due to IRB restrictions, demographic data or other metadata were not available for these samples.

#### Rhesus macaques

We obtained extracted DNA from rhesus macaque tissue samples (n=96) collected by the Cayo Biobank Research Unit in partnership with the University of Puerto Rico’s Caribbean Primate Research Center (CPRC) (95–99). Beginning in 2016, samples were collected from individuals living on the island of Cayo Santiago, an NIH-managed free-range colony of provisioned rhesus macaques. Specifically, as part of an ongoing population management plan designed by CPRC, select individuals were culled and tissues from all major organ systems were systematically harvested, stored in a fixative buffer, and frozen at -80C (IACUC #A400117). This data set consists of samples from six different tissue types: adrenal, heart, kidney, lung, liver, and spleen, with 16 samples from each type and samples coming from 23 unique individuals (Table S3). This dataset includes samples from 11 females and 12 males, ages 3.2 to 19.9 years old (mean 10.6 years old), collected from 2016–2019 (Table 1; Table S3). Genomic DNA was extracted using the Zymo *Quick*-DNA/RNA MagBead kit (Zymo Research #R2131) following the provided manufacturer’s protocols.

#### Geladas

We extracted DNA from whole blood from 68 geladas; 21 were female and 47 were male and all were considered adult (i.e., over 5 years old, the typical age of reproductive maturation in this species (100)) (Table 1). Gelada samples were collected as part of the Simien Mountains Gelada Research Project (SMGRP) which, since 2017, has carried out annual capture-and-release campaigns to collect morphometric data and whole blood samples from wild Ethiopian geladas. Samples were stored in liquid nitrogen upon collection, and at -80C for longer term storage. This research was conducted with approval from the Ethiopian Wildlife and Conservation Agency (EWCA), and the Institutional Animal Care and Use Committees (IACUCs) at the University of Washington (protocol 4416-01) and Arizona State University (20-1754 R) (see Chiou et al. for more details on sample collection (101)). Genomic DNA was extracted using the Qiagen DNeasy Blood & Tissue kits (Qiagen #69581) following the provided protocols.

#### Capuchins

Blood was collected from individuals in the captive capuchin monkey colony at Georgia State University in January 2023. Of the 28 capuchins, 19 were female and 9 were male with an average age of 19.4 years old (range 9–41 years old; Table 1). A trained veterinarian anesthetized the monkeys using 13 mg/kg Ketamine, delivered intramuscularly. Whole blood samples were collected in 6mL EDTA tubes, stored at 4°C, and shipped to Arizona State University where they were flash frozen into 0.5mL aliquots and stored at -80°C until used for analysis. DNA was extracted using the Qiagen DNeasy Blood & Tissue kits (Qiagen #69581) following the manufacturer’s protocols. Blood collection was approved by the GSU IACUC (protocol A20018).

### Overview of TMS library preparation

We used the Qubit dsDNA assay to determine the quantity of all extracted DNA. The samples were normalized to the desired input amount, and libraries were prepared using the NEBNext® Enzymatic Methyl-seq kit (P/N: E7120L) following a modified version of the manufacturer’s protocol that included 9 cycles of PCR for the final library amplification followed by a 0.65X bead cleanup. 71.4 ng of each resulting library was pooled and captured using the Human Methylome panel from Twist Biosciences following the manufacturer’s instructions (P/N: 105521). The final post-capture PCR reaction was split into 2 reactions per pool and cleaned with a 1X bead cleanup and then combined. Pool quality was assessed post-hybridization using the Agilent Bioanalyzer and quantified using a qPCR-based method with the KAPA Library Quantification Kit (P/N: KK4873) and the QuantStudio 12K instrument.

Prepared library pools were sequenced on the NovaSeq 6000 at the Vanderbilt Technologies for Advanced Genomics (VANTAGE) Core. We used 150 bp paired-end sequencing and targeted 30-50M paired-end reads per sample. Real Time Analysis Software (RTA) and NovaSeq Control Software (NCS) (1.8.0; Illumina) were used for base calling. MultiQC (v1.7; Illumina) was used for data quality assessments. For each sample, we applied the Illumina DRAGEN Methylation Pipeline v4.1.23 using the custom bed file from Twist Biosciences. The deliverables from DRAGEN consist of FASTQs, bams, cytosine reports (which include counts of methylated and unmethylated reads per CpG site), and methyl and mapping metric reports.

### TMS library preparation for experiments 1 & 2: Varying multiplexing strategies and input amounts

To determine whether TMS can be effectively multiplexed beyond the recommended 8-plex, we used 96 Tsimane samples to test four different multiplexing strategies during capture: 12-, 24-, 48-, and 96-plex. To test whether TMS is robust to DNA input amounts, we tested five input amounts: specifically, 25, 50, 100, 200, and 400 ng of sample were used as input into the EM-seq library prep. Here, we kept the plexing strategy constant (12-plex) and used three Tsimane samples, each represented three times within each pool and included three replicates of a control sample (HG01879 from the 1000 Genomes Project) (102).

### TMS library preparation for experiments 3 & 4: Optimizing capture efficiency and DNA shearing strategies

To optimize the capture efficiency of Twist target sites, we tested the use of two different annealing temperatures–65° and 68° C–along with the use of a methylation enhancer (ME)– produced by Twist Biosciences (Catalog #103557) consisting of Tris EDTA buffer to block the binding of off-target probes thereby improving on-target capture efficiency. The specific combinations we explored were: testing a 65°C annealing temperature with 0uL (n=192), 2uL (n=96), and 4uL (n=96) of ME and testing a 68°C annealing temperature with 0uL (n=96) and 2uL (n=192) of ME. These experiments were conducted with 96-plexed Tsimane samples (n=192), and using 200 ng of sample input.

Next, we tested the use of an enzymatic fragmentation method to replace the Covaris (LE220) mechanical shearing method to fragment the DNA. Mechanical shearing is known to decrease library quality through damage to DNA; however, previous use of enzymatic shearing methods have been shown to remove methyl groups from methylated CpG sites, thus biasing detected methylation levels. Here, we aim to test whether enzymatic fragmentation, which is not currently recommended by the TMS protocol, impacts site-level DNAm estimates through a comparison with samples prepared using mechanical shearing. We performed the optimized TMS with enzymatic shearing using 4uL of NEBNext UltraShear (NEB #M7634S/L) for 10 or 20 minutes. This experiment was conducted using 96-plexed samples from the Orang Asli (n=88) and using 200 ng of sample input.

### TMS and RRBS library preparation for experiments 7 and 8

To evaluate the efficacy of optimized TMS on three NHP species—macaques, geladas, and capuchins—we applied the 96-plex protocol design from experiment 1 with 200 ng input. To compare rhesus macaque TMS to RRBS, we generated libraries using 150 ng of DNA input in combination with 1ng of lambda phage DNA and 1uL of Msp1—a digestive enzyme which cuts CCGG nucleotide motifs. Next, using NEBNext Ultra II reagents, we performed end repair and adapter ligation to the DNA fragments produced by Msp1 digestion. We then performed bisulfite conversion on the fragments using the EZ-96 DNA Methylation Lightning MagPrep kit (Zymo Research #D5046) following the manufacturer directions. The samples were then PCR amplified for 16 cycles with unique dual indexed sequencing primers. We selected for 180-2000 bp fragments and sequenced on an Illumina NovaSeq S2 flow cell with 2x51bp sequencing (78,103).

### Low-level processing of TMS data

For experiments 1, 2, 7, and 8, we used a custom bioinformatics pipeline to process all FASTQ files into counts of methylated versus unmethylated cytosines at each CpG site. For experiments 3, 4, 5, and 6, we used Illumina’s Dynamic Read Analysis for GENomics (DRAGEN) pipeline (104) to process all FASTQ files into counts of methylated versus unmethylated cytosines at each CpG site. Importantly, both our custom pipeline and DRAGEN follow the same general steps and rely on the Bismark suite (105), making them highly comparable. We also processed 7 samples from experiment 4 using both methods to empirically confirm that our custom pipeline and the Illumina DRAGEN pipeline produced near identical results (Figure S13).

For our custom pipeline, we first trimmed adapters using Trimmomatic (version 0.39) (106) and TrimGalore (version 0.6.6) (107) for human and NHP samples, respectively. Following trimming, we used Bismark (version 0.24.0) (105) to map reads to each species’ respective genomes (hg38 for human, mmul10 for rhesus macaque, cimit for capuchin, and tgel1 for gelada). We retained only uniquely mapped reads and used the methylation extractor function within Bismark to extract counts of methylated versus unmethylated cytosines at each cytosine. These files were further filtered for CpG contexts only.

For all samples, run through either the custom or DRAGEN pipeline, we extracted two measures of data quality that are automatically calculated by Bismark: the percent of reads that mapped uniquely to the reference genome and the average methylation percentage for cytosines in a CHH context. The latter value serves as a commonly used estimate of the efficiency with which a given protocol converts unmethylated cytosines to thymine, because cytosines located outside of CpG contexts are extremely unlikely to be methylated in the mammalian genome (108,109). For experiments 1 and 2, we tested whether multiplexing strategy and input amount impacted mapping efficiency and percent CHH methylation using a one-way ANOVA test, followed by a pairwise t-test in the case of significance, with the ‘aov’ and ‘pairwise.t.test’ functions in the ‘stats’ R package (110).

For each study, we used the BSseq R package (111) to compile count matrices (derived from our custom pipeline or DRAGEN) across samples and to perform region, coverage, and missingness filtering. For experiments 1, 3, 4, 5, and 6 we used built-in functions in BSseq to filter for sites within the probes regions (+/- 200 bp) and for sites covered at >5X in >75% of samples. We made slight modifications to this filtering pipeline for other experiments. For experiment 2, where n=3 for each input amount, we relaxed our missingness filter to sites with at least one read observed in at least ⅔ samples. For experiment 7, which focused on NHP genomes for which the probe set coordinates (which are provided in hg38) are irrelevant, we did not perform region filtering. The number of sites analyzed for each experiment (reported in the main text and in Figure S2) therefore varies slightly depending on sample size, sequencing coverage, and other factors that impact which CpG sites passed our filters.

To confirm the fidelity of optimized TMS, we also checked whether CpGs captured by the protocol were distributed as expected throughout different genomic regions (e.g., promoters, enhancers) and that the average methylation levels in different genomic regions were as expected (e.g., quiescent regions being lowly methylated, repressed polycomb being highly methylated). To do so, we annotated each CpG site by whether it fell into a gene body, promoter, or non-genic region, and by chromatin state. We used hg38 gene body coordinates from Ensembl’s ‘biomaRt’ package in R, and we defined promoter regions as the 2000 bp region upstream of TSSs. We annotated CpGs as falling in chromatin states as defined by hg38 ChromHMM annotations from NIH’s Roadmap Epigenomics Project (50). We then counted the number of CpG sites that fell in each region (Figure 2C) and evaluated the median methylation across samples (Figure 2D).

### Quantifying capture efficiency

A subset of our experiments sought to understand and optimize two measures of efficiency of the hybrid capture step: 1) how many of the expected CpG sites from the probe set passed our filtering parameters and were thus analyzable and 2) how many of the reads we generated for a given sample were on-target and putatively captured by the probe set, rather than representing off-target randomly sequenced DNA fragments that do not contribute to analyzable data as they are often sparsely shared between samples. For #1, we used the bedtools (version 2.28.0) (112) intersect command to determine the proportion of CpG sites that are within +/- 200bp with at least 1 probe (using a bed file available on the Twist Biosciences website (https://www.twistbioscience.com/resources/data-files/twist-human-methylome-panel-target-bed-file). For #2, we used the bedtools function bamtobed to convert the mapped reads for each sample into a bed file; because we used a paired end sequencing strategy, each bed coordinate included a fragment start position from R1 and a fragment end position from R2. We then used the bedtools intersect command to determine the proportion of mapped read pairs that are within 200bp of at least 1 Twist probe.

### Comparing DNA methylation measurements between TMS, the EPIC array, and WGBS

We used our filtered BSSeq object from experiment 5 to compare to data from the EPIC array generated for 55 paired human samples (average number of CpG sites measured with EPIC = 936,280; average call rate = 0.999). We downloaded the EPIC CpG coordinates from the Illumina website and merged with the TMS CpG locations, resulting in a shared dataset of 682,295 CpG sites passing filters and common to both technologies. We then performed two analyses to understand consistency. First, we calculated the average per-site methylation level across all samples included in the TMS or EPIC array datasets, respectively. We then ran a linear model testing the relationship between the two sets of average methylation levels using the ‘lm’ function in the ‘stats’ package in R. Second, we used the ‘lm’ function to estimate the R^2^ value comparing per-site methylation levels for estimates derived from each technology for a given individual (i.e., not averaged across the dataset). This resulted in a distribution of 55 R^2^ values. Because all humans share canonical methylation patterns, we also compared this distribution to a distribution of 55 R^2^ values derived from the same analysis after sample identity was permuted. We used the ‘t.test’ function in the ‘stats’ package in R to ask whether these distributions were significantly different.

We used a very similar strategy to compare ∼30x WGBS data generated for six paired Tsimane samples with TMS data generated from experiment 1 (96-plex, 200 ng input). First, we performed low level processing of the WGBS data using Illumina’s DRAGEN pipeline and merged this with our filtered TMS data, resulting in 3,078,771 CpG sites common to both datasets. We calculated the average methylation level across samples reported for each site and technology and ran a linear model using the ‘lm’ function in the ‘stats’ package in R to calculate the R^2^ value. We did not compare individual-based R^2^ values to permuted values for this experiment, given the small number of individuals.

### Understanding TMS performance in NHP species and comparing DNA methylation measurements between TMS and RRBS

To estimate the number of CpG sites that we expected to recover when applying the human probe set to each NHP species, we converted the probe bed file to a FASTA file using the bedtools command ‘getfasta’ (112) and the hg38 reference genome. We then used Bismark to map the FASTA file to each non-human primate’s respective genome. From the mapped bam file, we used the ‘bamToBed’ function in bedtools to extract coordinates for the mapped probes and to add a +/- 200 bp offset. Finally, we applied the ‘getfasta’ function in bedtools to extract the sequence for the mapped regions (plus the 200bp buffer) from the non-human primate genome and to count the number of CpG sites in this region set.

Similar to the comparisons between TMS and the EPIC array, we used paired RRBS data for the 96 rhesus macaque samples to directly compare methylation data generated using TMS versus RRBS. To do so, we processed the RRBS data using the same custom pipeline and filtering parameters described for TMS data, with the only modification being that we used the ‘—rrbs’ parameter in Trim Galore to remove unmethylated cytosines artificially introduced during library preparation from the 3’ end of fragments. We merged the filtered TMS and RRBS datasets, resulting in 721,766 CpG sites common to both technologies. As described for the TMS-EPIC array comparison, we then 1) calculated the average per-site methylation level across all samples included in each dataset and compared these vectors using linear models and 2) estimated the R^2^ value for methylation level estimates derived from each technology for a given individual, and used a t-test to compare this distribution to a distribution for the same analysis where sample identity was permuted (Figure S14).

### Testing for age-associated changes in DNA methylation in the Tsimane and Orang Asli

We tested the extent to which DNA methylation varied with age using data from two experiments: 1) 96-plex, 200 ng input data from experiment 1 for the Tsimane of Bolivia and 2) 96-plex, 200 ng input, enzymatically sheared data from experiment 4 for the Orang Asli of Peninsular Malaysia. Subsistence-level groups generally show little age-related decline in cardiometabolic health (113), but it remains unknown if their relative lack of age-related chronic disease extends to molecular phenotypes because these populations are chronically understudied and underrepresented, especially in genomics. To evaluate age-effects on DNA methylation levels, we focused on sites that passed the filtering parameters described above and that were variably methylated (median methylation >10% and <90%). This resulted in 936,547 and 1,001,011 testable sites in the Tsimane and Orang Asli, respectively. We used the ‘betabin’ function from the R ‘aod’ package (114) to test for age effects on the proportion of methylated / total read counts at each CpG site in each population, controlling for self-reported sex, batch, and cell type composition (proportion of neutrophils and lymphocytes, including B, CD4-T, and CD8-T cells). We extracted the p-value for the age effect for each tested site, and corrected for multiple hypothesis testing using the Benjaminin-Hochberg false discovery rate approach implemented in the ‘p.adjust’ function in R.

Cell type composition was estimated via deconvolution in the R package ‘EpiDISH’ (115), using the ‘centDHSbloodDMC.m’ reference and the ‘RPC’ method. We focused on the cell populations referenced above because they matched empirically derived estimates available for the same samples for the Orang Asli (R^2^>0.5, p<0.05). Specifically, information about the relative proportion of lymphocytes, neutrophils, eosinophils, monocytes, and basophils was obtained for each Orang Asli sample from a 5-part white blood cell differential via the HemoCue WBC DIFF system and compared to deconvolution estimates.

We compared the age-related effects we found in Tsimane and Orang Asli to those reported by Johansson et al. (2013) (70), which quantified DNA methylation throughout aging in a Swedish cohort using the 450K array. As Johansson et al. provided age estimates by Illumina ID and chromosome/position (i.e., site), we used UCSC’s Genome Browser LiftOver to convert genomic coordinates to hg38. We then were able to combine this dataset with our TMS data based on genomic coordinate position. This allowed us to test for correlations of age effect sizes amongst CpGs with data in all three datasets (Johansson, TMS from Tsimane, and TMS from Orang Asli). To understand how age effects varied across regions of the genome with distinct functions, we annotated each CpG site by the chromatin states in which it fell using ChromHMM annotations from NIH’s Roadmap Epigenomics Project (50). We then asked—for each population separately—whether CpGs significantly hypermethylated with age (FDR < 10%) were enriched in each chromatin state compared to all other sites and age-associated hypermethylated sites that did not fall into that region using the ‘fisher.test’ function in R. We performed this test for all regions and did the same for hypomethylated sites. All statistical analyses were performed using R version 4.2.2.

## DATA AND CODE AVAILABILITY

All NHP data generated as part of this study has been deposited in NCBI’s Sequence Read Archive under accession number PRJNA1156067.

The human genomic data generated as part of this study comes from Indigenous participants from the Tsimane Health and Life History Project (THLHP) and the Orang Asli Health and Lifeways Project (OA HeLP). Both THLHP and OA HeLP’s highest priority is the minimization of risk to study participants. Both projects adhere to the “CARE Principles for Indigenous Data Governance” (Collective Benefit, Authority to Control, Responsibility, and Ethics) and are also committed to the “FAIR Guiding Principles for scientific data management and stewardship” (Findable, Accessible, Interoperable, Reusable). To adhere to these principles while minimizing risks, genomic data from both projects are available via restricted access. For OA HeLP, these requests can be made via dbGap (accession number TBD). For THLHP, these requests can be made via email following the instructions provided here: https://tsimane.anth.ucsb.edu/data.html. In both cases, requests for de-identified genomic data will take the form of an application that details the exact uses of the data and the research questions to be addressed, procedures that will be employed for data security and privacy, potential benefits to the study communities, and procedures for assessing and minimizing stigmatizing interpretations of the research results. Both projects are committed to open science and the leadership is available to assist interested investigators in preparing data access requests.

All scripts used to perform the analyses described here are available at “https://github.com/alongtin15/TMS-Cost-effective-solutions-for-high-throughput-enzymatic-DNA-methylation-sequencing.”

## Supporting information

Complete SI Materials

Longtin 2024 TableS1

Longtin 2024 TableS16

Longtin 2024 Table S17

## ACKNOWLEDGMENTS

First, we thank the Tsimane and Orang Asli study participants and communities for their involvement and support. We also thank all members of the research teams associated with the Tsimane Health and Life Histories Project and the Orang Asli Health and Lifeways Project. Second, we thank the members of the Lea Lab at Vanderbilt University, the SMack Lab at Arizona State University, and the Vanderbilt Technologies for Advanced Genomics (VANTAGE) team for their feedback, expertise, and support in completing this work. Third, we are grateful to the research infrastructure provided by Vanderbilt University’s Advanced Computing Center for Research and Education and Arizona State University’s Sol Computing clusters. Fourth, we thank Dr. Rex Howard and Dr. Michael Hart, the GSU veterinarians, for their assistance in collecting the capuchin blood samples. Finally, we acknowledge funding from the National Institute of General Medical Sciences (R35-GM147267), National Institute on Ageing (R01AG054442 and R61AG078529), the National Science Foundation (BCS-2142090), the Canadian Institute for Advanced Research (Azrieli Global Scholars Program), the French National Research Agency under the Investments for the Future (Investissements d’Avenir) programme (ANR-17-EURE-0010), the Kinship Foundation, (Searle Scholars Program), and the Pew Charitable Trusts (Pew Biomedical Scholars Program).

## SUPPLEMENTAL INFORMATION

Supplemental information provided in a separate document.

