## Supplementary material for "Cost-effective solutions for high-throughput enzymatic DNA methylation sequencing": Complete SI Materials

### SUPPLEMENTAL METHODS

#### Study populations included in epigenomic aging analyses

The Tsimane Health and Life History Project (THLHP) has conducted long-term anthropological and biomedical research with the Tsimane people for several decades (1). The Tsimane are an Amerindian population from the Bolivian Piedmont and maintain a subsistence-level lifestyle supported by slash‐and‐burn horticulture, fishing, hunting, and foraging (1). They experience extremely low rates of non-communicable, lifestyle-associated diseases (2). The THLHP regularly monitors the demography, behavior, and health of thousands of Tsimane individuals, including through regular medical exams that include the collection of whole blood samples used for this study.

We extracted DNA from 192 venous whole blood (WB) samples collected between the years of 2010–2021 as part of the Tsimane Health and Life History Project (THLHP). Samples were frozen in liquid nitrogen, transferred on dry ice to Arizona State University, and stored at -80ºC prior to analysis. Informed consent was collected at three levels: by the individual, by the community, and by the Tsimane Gran Consejo (Tsimane governing body). All study protocols were approved by the Institutional Review Boards of the University of New Mexico (# 07-157), the University of California Santa Barbara (#3-21-0652), and Universidad Mayor San Simon, Cochabomba.

The Orang Asli are the Indigenous peoples of Peninsular Malaysia, comprising <1% (~210,000 people) of the country's population. They are typically divided in 19 distinct ethnolinguistic groups and three broader sub-groups: the Negrito (traditionally nomadic hunter-gatherers), the Senoi (traditionally horticulturalists), and the Proto-Malay (traditionally mixed subsistence practitioners). The Orang Asli show genomic evidence of a long history in their ancestral environments (3), and their ecology and biometric phenotypes have been well-studied by anthropologists for several decades (4,5). Over the last few decades, Orang Asli in Malaysia have undergone rapid lifestyle change due to government-promoted assimilation, deforestation driven by natural resource extraction, and the expansion of built infrastructure (6). The Orang Asli Health and Lifeways Project (OA HeLP) has worked with >1000 individuals across this environmental gradient to collect data on behavior, culture, genetics, and physiology to understand how rapid lifestyle change impacts health (7).

Orang Asli samples were collected in May-July of 2023. Samples were frozen in liquid nitrogen upon collection, exported to Vanderbilt University in a dry shipper, and subsequently stored at -80ºC. Similar to the THLHP, informed consent was collected by first describing the project to the community as a whole and seeking the permission of community leaders, and subsequently through individual-specific review of the protocol.

**Parameters used for low level data processing of TMS data**

For experiments 1, 2, 7, and 8, we used a custom bioinformatics pipeline to process all FASTQ files into counts of methylated versus unmethylated cytosines at each CpG site. We trimmed the FASTQ files from experiments 1 and 2 using Trimmomatic (ILLUMINACLIP:TruSeq3-PE.fa: 2:30:10:8:trueHEADCROP:3 TRAILING:10 MINLEN:25) (version 0.39) (8) and we trimmed the FASTQ files from experiments 7 and 8 using TrimGalore (--paired, -j 8) (version 0.6.6) (9). Following trimming, we processed all files using Bismark (version 0.24.0) (10) for mapping (hg38 for human, mmul10 for macaque, cimit for capuchin, and tgel1 for gelada) (--score_min L,0,-0.6) and methylation extraction (bismark_methylation_extractor: -p, --bedGraph, --comprehensive; coverage2cytosine: --merge_CpG). For experiments 3, 4, 5, and 6, we downloaded cytosine reports (the equivalent files produced from Illumina's DRAGEN pipeline, which relies on the Bismark suite (10)) from Ilumina's BaseSpace CLI interactive portal.

This Illumina's DRAGEN pipeline as well as the in-house commands described above generate 1) a cov.gz file used for downstream analyses and 2) multiple report files containing information such as mapping efficiency and average CHH methylation level (which were extracted and summarized in the main text). We used the .cov.gz files and combined them within a given experiment using the BSseq package in R (11) to generate an RDS objects using the read.bismark command (files = list.files, verbose = TRUE, strandCollapse = TRUE, backEnd = "HDF5Array"). For each experiment, this object was filtered (using built in functions in BSseq) to include CpG sites within the Twist probe set, that were covered at >5x on average across samples, and that had data for >75% of samples.

**Estimating the number of reads needed to achieve a given coverage**

We used data from experiment 4, which represents our "best" protocol (96-plex, 200ng input, 65ºC annealing, no methylation enhancer, enzymatic shearing), to understand a critical aspect of experimental design—how many reads one would need to generate to achieve a given average coverage per CpG site. To do so, we used the 'view' function (-@ 20 -bh -s) in SAMtools (12) to subsample the uniquely mapped reads resulting from the enzymatic fragmentation experiment for each sample (specifically subsampling to include 25%, 50%, and

75% of the mapped reads in each bam file). We then reran our sample processing pipeline starting with the bam/mapped reads file (see previous section) and calculated the average coverage per CpG site across all sites represented by each original file or subsampled file. We then compared the number of uniquely mapped reads to this value. In general, we observe a 1:2 relationship between the number of mapped, paired end reads (in millions) and average coverage, such that ~25M mapped paired end reads translates to ~50x average coverage per CpG site. The results of this analysis can be found in Figure S5.

### SUPPLEMENTAL TABLES

**Table S1: Read depth and metadata per human sample, broken down by experiment and condition.** See table S16 for read depth by non-human primate sample. Table is provided in a separate XLSX file “Longtin 2024_TableS1.xlsx.”

**Table S2: Comparison of mapping efficiency with varying DNA input amounts.** P-values generated from pairwise t-tests comparing the percentage of reads that were uniquely mapped to the human genome from sequencing data generated using the TMS protocol with varying amounts of input DNA. * represents a significant ($p < 0.05$) difference in mapping efficiency between conditions.

| | 25 ng | 50 ng | 100 ng | 200 ng | 400 ng |
|---|---|---|---|---|---|
| **25 ng** | - | - | - | - | - |
| **50 ng** | 0.1453 | - | - | - | - |
| **100 ng** | 0.0044* | 0.0518 | - | - | - |
| **200 ng** | 0.0012* | 0.0066* | 0.2363 | - | - |
| **400 ng** | 0.0011* | 0.0043* | 0.1209 | 0.5728 | - |

**Table S3: Comparison of CHH methylation with varying plexing strategies.** P-values generated from pairwise t-tests comparing the percentage of cytosines in a CHH context marked as methylated (an estimate of conversion efficiency). * represents a significant ($p < 0.05$) difference in percent CHH methylation between conditions.

| | 12-plex | 24-plex | 48-plex | 96-plex |
|---|---|---|---|---|
| **12-plex** | - | - | - | - |
| **24-plex** | 1 | - | - | - |
| **48-plex** | 1 | 1 | - | - |
| **96-plex** | $< 2 \times 10^{-16}$ * | $< 2 \times 10^{-16}$ * | $< 2 \times 10^{-16}$ * | - |

**Table S4: Comparison of CHH methylation with varying input amounts.** P-values generated from pairwise t-tests comparing the percentage of cytosines in a CHH context marked as methylated (an estimate of conversion efficiency). * represents a significant ($p < 0.05$) difference in percent CHH methylation between conditions.

| | 25 ng | 50 ng | 100 ng | 200 ng | 400 ng |
|---|---|---|---|---|---|

| | | | | | |
|---|---|---|---|---|---|
| **25 ng** | - | - | - | - | - |
| **50 ng** | $< 2 \times 10^{-16}$ * | | - | - | - |
| **100 ng** | $< 2 \times 10^{-16}$ * | 1 | - | - | - |
| **200 ng** | $< 2 \times 10^{-16}$ * | 1 | 1 | - | - |
| **400 ng** | 1 | $< 2 \times 10^{-16}$ * | $< 2 \times 10^{-16}$ * | $< 2 \times 10^{-16}$ * | - |

**Table S5: Number of captured on-target sites and average site-based coverage for each plexing strategy.** Sites are filtered for those within the Twist probe set and covered at >5x coverage in >75% of samples.

| Plexing Strategy | Number of Sites | Average Coverage |
|---|---|---|
| 12-plex | 4,970,970 | 89.37811 |
| 24-plex | 4,049,084 | 28.50994 |
| 48-plex | 4,083,989 | 23.0657 |
| 96-plex | 3,683,988 | 21.46068 |

**Table S6: Number of captured on-target sites and average site-based coverage for each input amount.** Sites are filtered for those within the Twist probe set and covered at >5x coverage in >75% of samples.

| Input Amount (ng) | Number of Sites | Average Coverage |
|---|---|---|
| 25 | 3,943,058 | 23.01141 |
| 50 | 4,089,839 | 25.41013 |
| 100 | 4,189,354 | 33.87878 |
| 200 | 4,030,859 | 34.49294 |
| 400 | 4,006,596 | 37.19262 |

**Table S7: Percent of probes represented for each plexing strategy.** The percentage of Twist target probes (n= 551,803) covered by at least one read for each plexing strategy.

| Plexing Strategy | Percent of Probes Represented |
|---|---|

| | |
|---|---|
| 12-plex | 96.83% |
| 24-plex | 95.93% |
| 48-plex | 96.25% |
| 96-plex | 96.68% |

**Table S8: Percent of probes represented for each input amount.** The percentage of Twist target probes (n= 551,803) covered by at least one read for each input amount.

| Input Amount (ng) | Percent of Probes Represented |
|---|---|
| 25 | 92.11% |
| 50 | 92.27% |
| 100 | 92.41% |
| 200 | 92.11% |
| 400 | 92.06% |

**Table S9: Correlation in average methylation at each site between plexing strategies.** $R^2$ values generated using linear modeling to compare average site-level methylation between plexing experiments. Average site-level methylation was calculated by averaging the percent methylation for each site across all samples within a given plexing strategy and comparing these with average site-level methylation within shared sites in an alternate plexing strategy. All sites were filtered for >5X coverage in >75% of samples.

| | 12-plex | 24-plex | 48-plex | 96-plex |
|---|---|---|---|---|
| **12-plex** | 1 | - | - | - |
| **24-plex** | 0.9986 | 1 | - | - |
| **48-plex** | 0.9987 | 0.9989 | 1 | - |
| **96-plex** | 0.9988 | 0.9991 | 0.9994 | 1 |

**Table S10: Correlation in average methylation at each site between input amounts.** $R^2$ values generated using linear modeling to compare average site-level methylation between input amount experiments. Average site-level methylation was calculated by averaging the percent methylation for each site across all samples within a given input amount experiment and comparing with average site-level methylation within shared sites in an alternate input amount experiment. All sites were filtered for >5X coverage in >75% of samples.

| | 25 ng | 50 ng | 100 ng | 200 ng | 400 ng |
|---|---|---|---|---|---|
| **25 ng** | 1 | - | - | - | - |
| **50 ng** | 0.9902 | 1 | - | - | - |
| **100 ng** | 0.9911 | 0.9921 | 1 | - | - |
| **200 ng** | 0.9908 | 0.9919 | 0.9929 | 1 | - |
| **400 ng** | 0.9909 | 0.9920 | 0.9930 | 0.9928 | 1 |

**Table S11: Comparison of percent off-target reads with varying protocol modifications.** P-values generated from pairwise t-tests comparing the percentage of the total reads that were not associated with a Twist target probe (within +/- 200 bp) for each capture efficiency experiment. 65C/68C refers to annealing temperature and 0uL/2uL/4uL ME refers to volume of methylation enhancer. * represents a significant ($p < 0.05$) difference in the percent of probes captured between conditions.

| | 65C, 0uL ME | 65C, 2uL ME | 65C, 4uL ME | 68C, 0uL ME | 68C, 2uL ME |
|---|---|---|---|---|---|
| **65C, 0uL ME** | - | - | - | - | - |
| **65C, 2uL ME** | $6.8 \times 10^{-5}$* | - | - | - | - |
| **65C, 4uL ME** | $1.0 \times 10^{-6}$* | $1.6 \times 10^{-9}$* | - | - | - |
| **68C, 0uL ME** | $2.8 \times 10^{-15}$* | $1.2 \times 10^{-13}$* | $2 \times 10^{-4}$* | - | - |
| **68C, 2uL ME** | $< 2 \times 10^{-16}$* | $< 2 \times 10^{-16}$* | $5.5 \times 10^{-14}$* | $5.1 \times 10^{-5}$* | - |

**Table S12: Comparison of probe capture with varying protocol modifications.** P-values generated from pairwise t-tests comparing the percentage of Twist target probes covered by at least one read for each capture efficiency experiment. 65C/68C refers to annealing temperature and 0uL/2uL/4uL ME refers to volume of methylation enhancer. * represents a significant ($p < 0.05$) difference in the percent of probes captured between conditions.

| | 65C, 0uL ME | 65C, 2uL ME | 65C, 4uL ME | 68C, 0uL ME | 68C, 2uL ME |
|---|---|---|---|---|---|
| **65C, 0uL ME** | - | - | - | - | - |
| **65C, 2uL ME** | 0.206 | - | - | - | - |
| **65C, 4uL ME** | 0.479 | 0.098 | - | - | - |

| | | | | | |
|---|---|---|---|---|---|
| **68C, 0uL ME** | $3.1 \times 10^{-13}$ * | $1.5 \times 10^{-7}$ * | $1.3 \times 10^{-12}$ * | - | - |
| **68C, 2uL ME** | $4 \times 10^{-13}$ * | $5.8 \times 10^{-6}$ * | $1.4 \times 10^{-11}$ * | 0.098 | - |

**Table S13: Number of captured on-target sites and average site-based coverage for each capture efficiency experiment.** Sites are filtered for those which are covered at >5x coverage in >75% of samples.

| Experiment | Number of Sites | Average Coverage |
|---|---|---|
| 65C, 0uL ME | 4,199,890 | 23.02 |
| 65C, 2uL ME | 4,297,931 | 21.85 |
| 65C, 4uL ME | 4,453,577 | 29.17 |
| 68C, 0uL ME | 3,422,658 | 25.83 |
| 68C, 2uL ME | 3,885,350 | 38.09 |

**Table S14: Correlation in methylation across DNA shearing methods.** $R^2$ values generated using linear modeling to compare site-specific methylation for 3 samples, each processed with 3 different shearing methods- mechanical, enzymatic for 10 minutes, and enzymatic for 20 minutes. All sites were filtered for >5X coverage in >75% of samples.

| Sample | 10 min vs mechanical ($R^2$) | 20 min vs mechanical ($R^2$) | 10 min vs 20 min ($R^2$) |
|---|---|---|---|
| 1 | 0.9735 | 0.9744 | 0.9866 |
| 2 | 0.967 | 0.9663 | 0.9869 |
| 3 | 0.939 | 0.9739 | 0.987 |

**Table S15: Rhesus macaque multi-tissue dataset.** Age, sex, and tissue types for each individual in the rhesus macaque multi-tissue dataset, used to assess the function of TMS in a NHP with a direct comparison to RRBS data generated from these same samples (see also Figure S8).

| Individual | Age (years) | Sex | Tissue type(s) | Number of Tissues Used |
|---|---|---|---|---|
| 22H | 19.9 | F | Adrenal, kidney, lung, spleen, liver | 5 |
| 6R9 | 3.2 | M | Adrenal, heart, kidney, lung, spleen | 5 |
| 5J3 | 9.2 | M | Kidney, lung, spleen, liver | 4 |

| | | | | |
|---|---|---|---|---|
| 8K7 | 6.2 | F | Adrenal, heart, kidney, lung, spleen, liver | 6 |
| 5H7 | 8.2 | M | Heart, liver | 2 |
| 5J6 | 7.1 | F | Adrenal, heart, kidney, lung, spleen, liver | 6 |
| 56K | 19.9 | M | Adrenal, heart, kidney, lung, spleen | 5 |
| 2M4 | 7.2 | M | Adrenal, heart, kidney, lung, spleen, liver | 6 |
| 2K0 | 8.1 | M | Adrenal, heart, kidney, lung, spleen, liver | 6 |
| 42T | 16.0 | M | Adrenal, heart, kidney, lung, spleen, liver | 6 |
| 91O | 15.9 | M | Adrenal, heart, kidney, spleen | 4 |
| 80S | 14.1 | M | Adrenal, kidney, lung, spleen | 4 |
| 7F0 | 11.2 | F | Adrenal, heart, kidney, lung, spleen, liver | 6 |
| 4Q4 | 5.2 | F | Adrenal, heart, kidney, lung, spleen, liver | 6 |
| 3T9 | 4.4 | F | Adrenal, heart, kidney, lung, spleen | 5 |
| 8B3 | 13.1 | M | Adrenal, heart, kidney, lung, spleen | 5 |
| 4C6 | 10.8 | M | Adrenal, heart, kidney, lung, spleen | 5 |
| 6D9 | 13.1 | F | Heart, liver | 2 |
| 6B4 | 11.2 | F | Liver | 1 |
| 3O3 | 6.2 | M | Liver | 1 |
| 51V | 15.1 | F | Liver | 1 |
| 3F7 | 11.2 | F | Heart, lung, liver | 3 |
| 5M5 | 7.2 | F | Liver | 1 |

**Table S16: Read depth by NHP species.** Table is provided in a separate XLSX file "Longtin 2024_TableS16.xlsx."

**Table S17: Age-associated sites, their populations, annotations, p-values, and direction of methylation change with age.** Table is provided in a separate XLSX file "Longtin 2024_TableS17.xlsx."

### SUPPLEMENTAL FIGURES

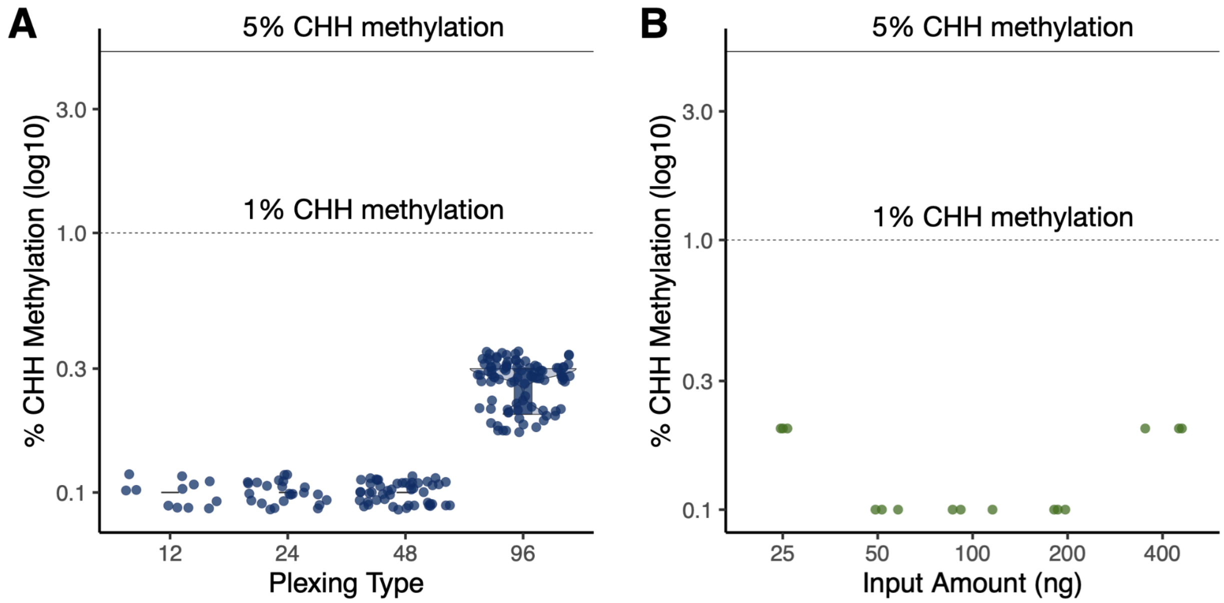


**Figure S1: Comparison of CHH methylation across experiments.** Percentage of cytosines in a CHH context marked as methylated (an estimate of conversion efficiency) for varying A) plexing strategies, and B) input amounts. The dashed line refers to 1% CHH methylation and the solid line refers to 5% CHH methylation, a common cut off indicative of high levels of unmethylated cytosine conversion.

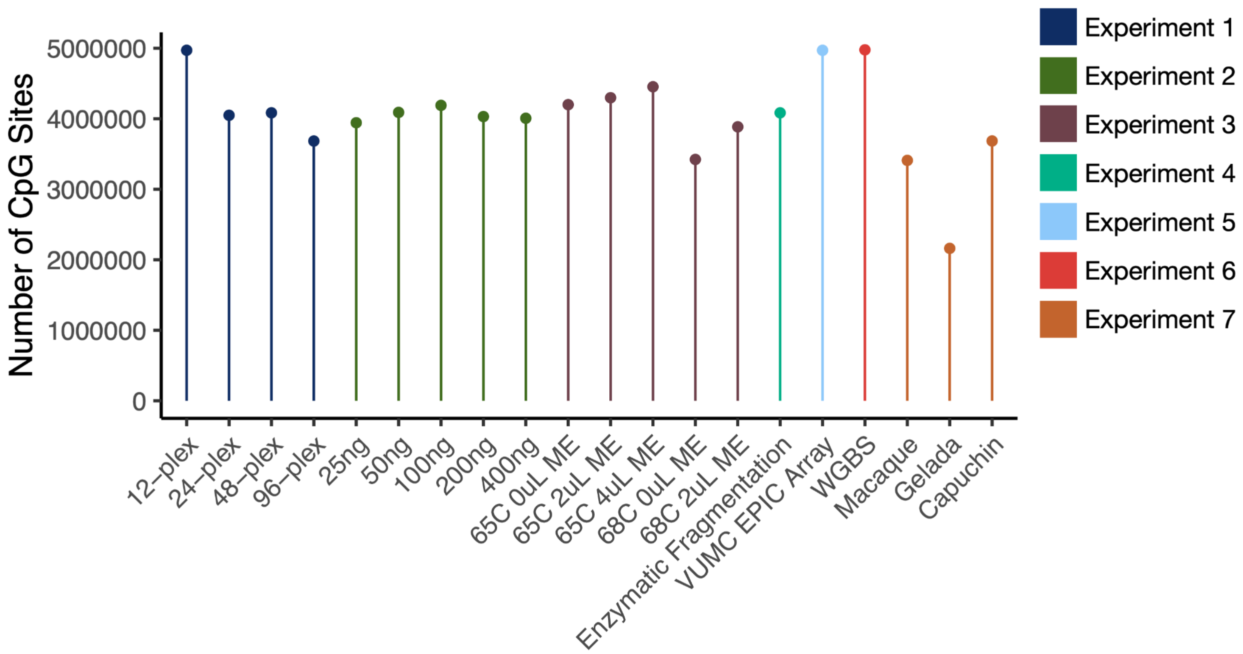


**Figure S2: Number of on-target CpG sites represented in each experiment.** Number of CpG sites within 200bp of target probes after filtering for >5X coverage in more than 75% of samples by experiment. Colors are representative of each experiment which are defined in Figure 1C.

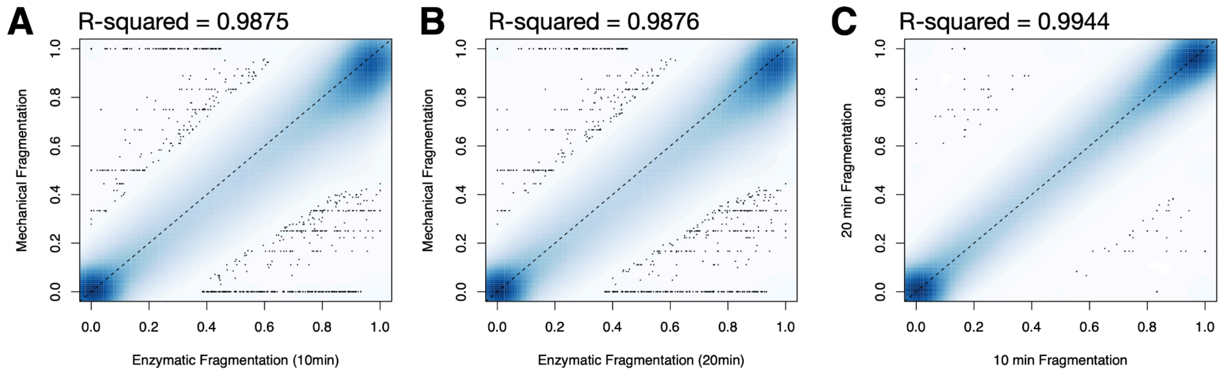


**Figure S4: Correlation in average site-level methylation for varying fragmentation methods.** Site-level methylation averaged across 3 samples processed using mechanical fragmentation, enzymatic fragmentation for 10 minutes, and enzymatic fragmentation of 20 minutes. Each point represents a site measured across both shearing methods and $R^2$ values were generated using linear modeling.

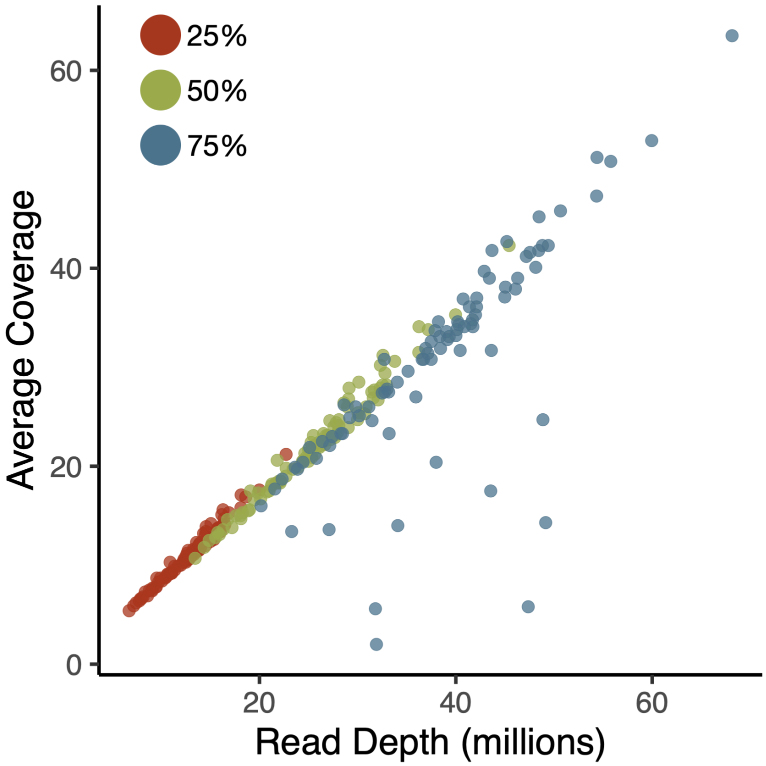


**Figure S5: Average coverage for on-target sites when mapped read files are subset to varying degrees.** We subset the mapped read files for each sample (n=88) included in our enzymatic fragmentation experiment (experiment 4) to include a random subset of 25, 50, or 75% of the total reads. We calculated the average coverage for on-target sites (y-axis) and observed a linear relationship between coverage and the number of subset reads (x -axis), which is useful for estimating what sequencing depth per sample will be needed to obtain various degrees of coverage.

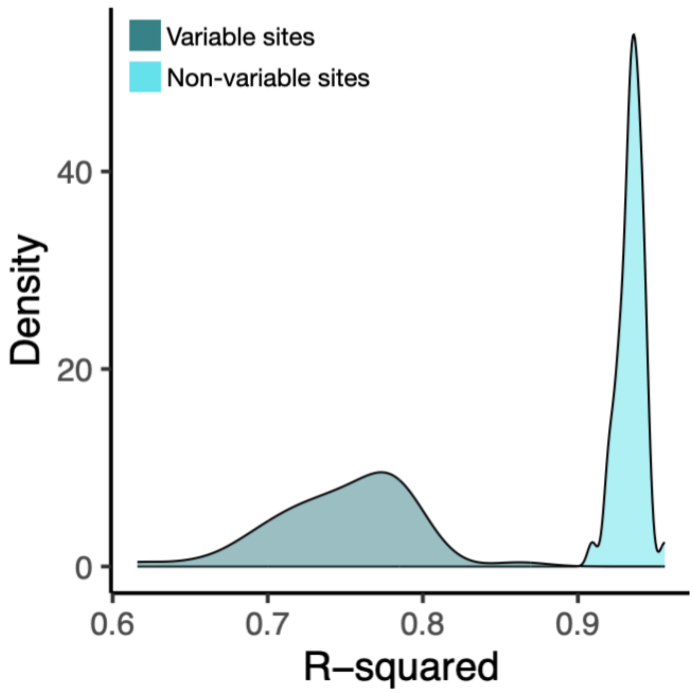


**Figure S6: Correlation in site-level methylation between TMS and EPIC array data after the permutation of sample identifiers.** For samples processed using both TMS and the EPIC array, we assessed the correlation in site-level methylation for variable sites (methylation >0.1 and <0.9) and non-variable sites (methylation < 0.1 or >0.9) after permuting sample ID randomly. $R^2$ values were generated using linear modeling.

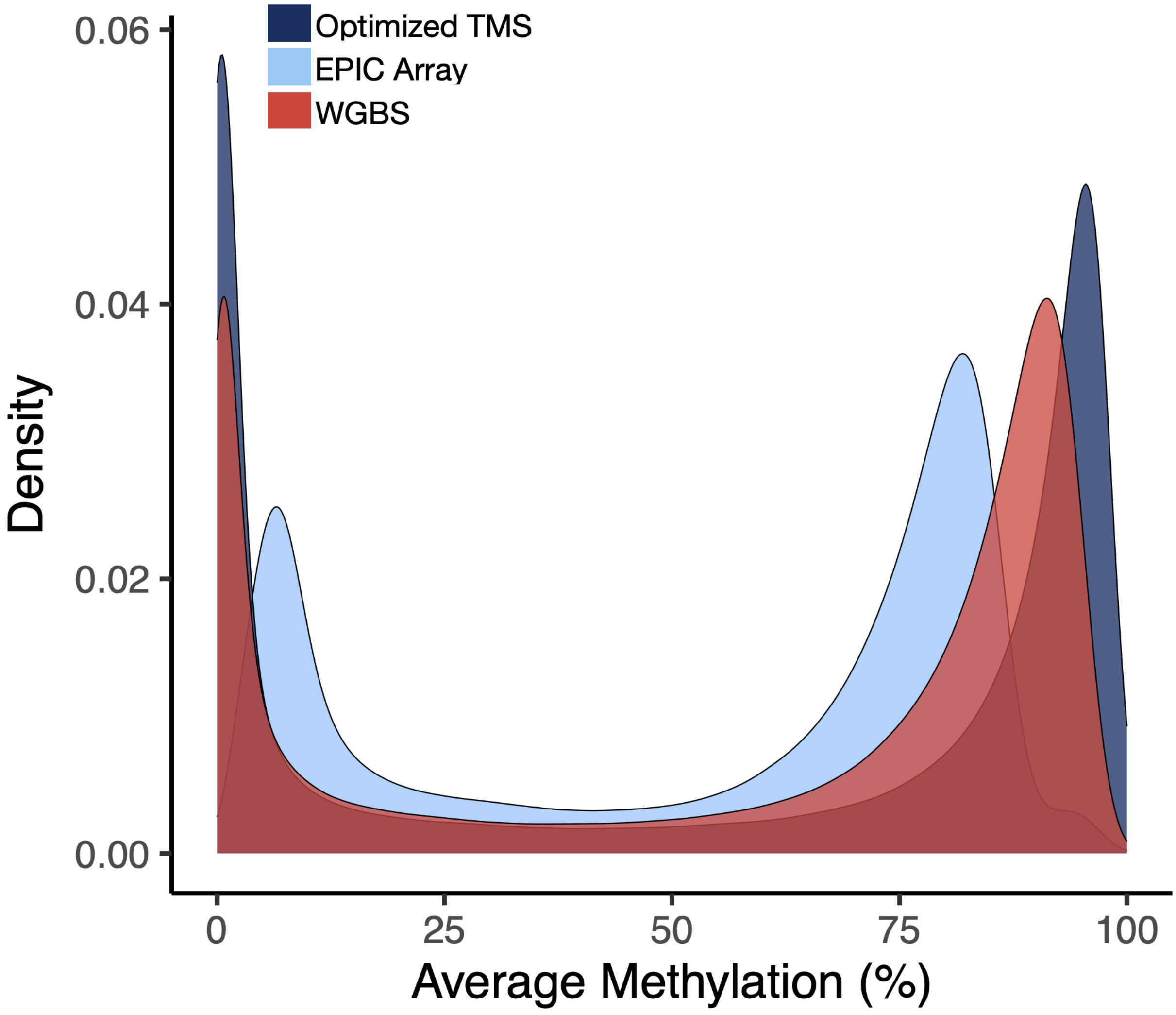


**Figure S7: Density plot showing the average methylation of a site (i.e., across samples) for filtered (>5X coverage in >75% of sites) sites captured between the three technologies (726,597 EPIC Array sites; 4,990,351 TMS sites; and 5,000,659 WGBS sites).** Sites were not matched between the three technologies. Additionally, WGBS samples are not paired with the TMS and EPIC Array samples. TMS and EPIC Array samples are paired.

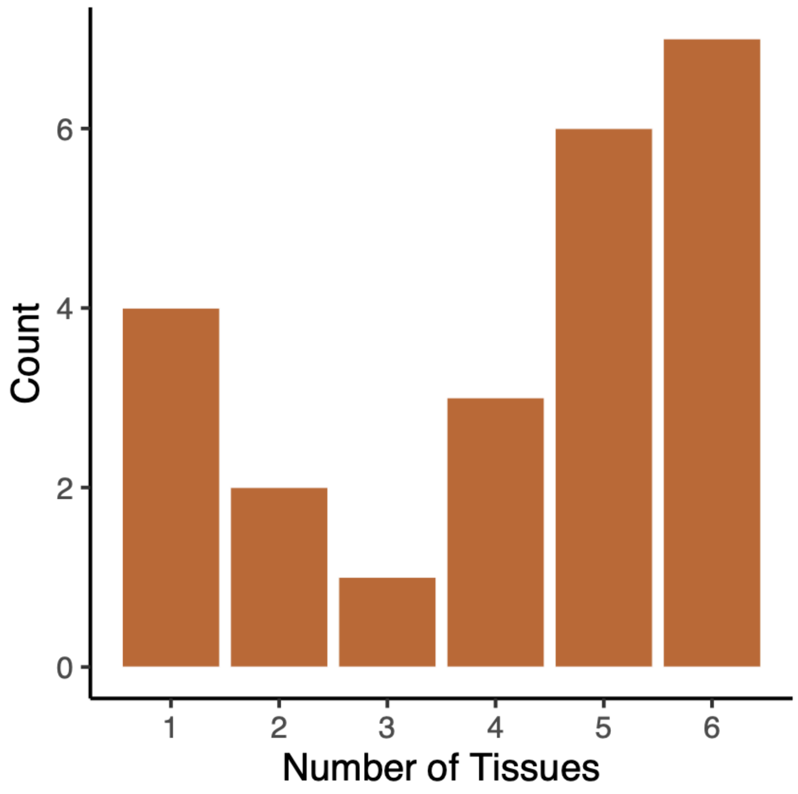


**Figure S8: Number of individuals from which different numbers of tissues were included in the rhesus macaque multi-tissue dataset.** The majority of individuals had 4+ tissues represented in the dataset.

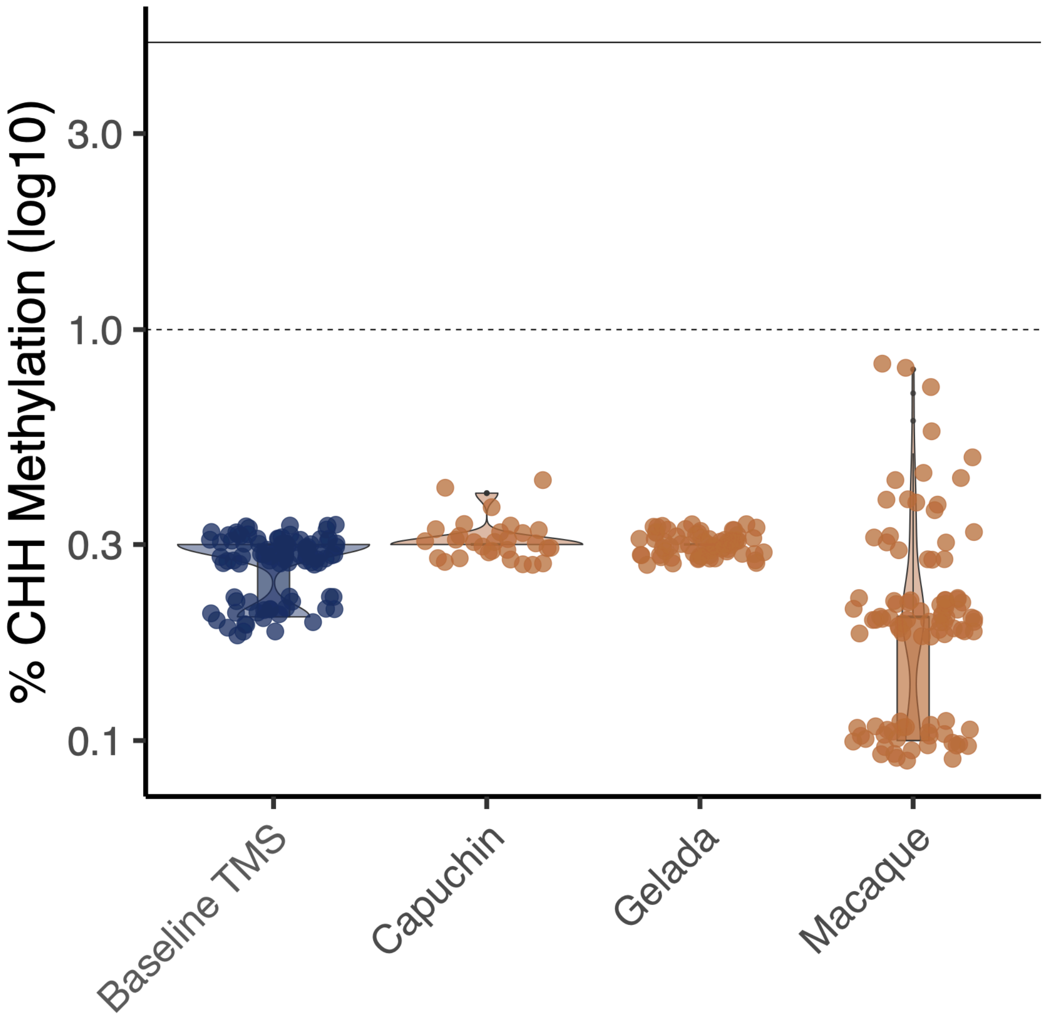


**Figure S9: Comparison of CHH methylation across experiments testing optimized TMS in three NHP species.** Percentage of cytosines in a CHH context marked as methylated (an estimate of conversion efficiency) following optimized TMS using genomic DNA from capuchins, geladas, and macaques. The dashed line refers to 1% CHH methylation and the solid line refers to 5% CHH methylation, a common cut off indicative of high levels of cytosine conversion.

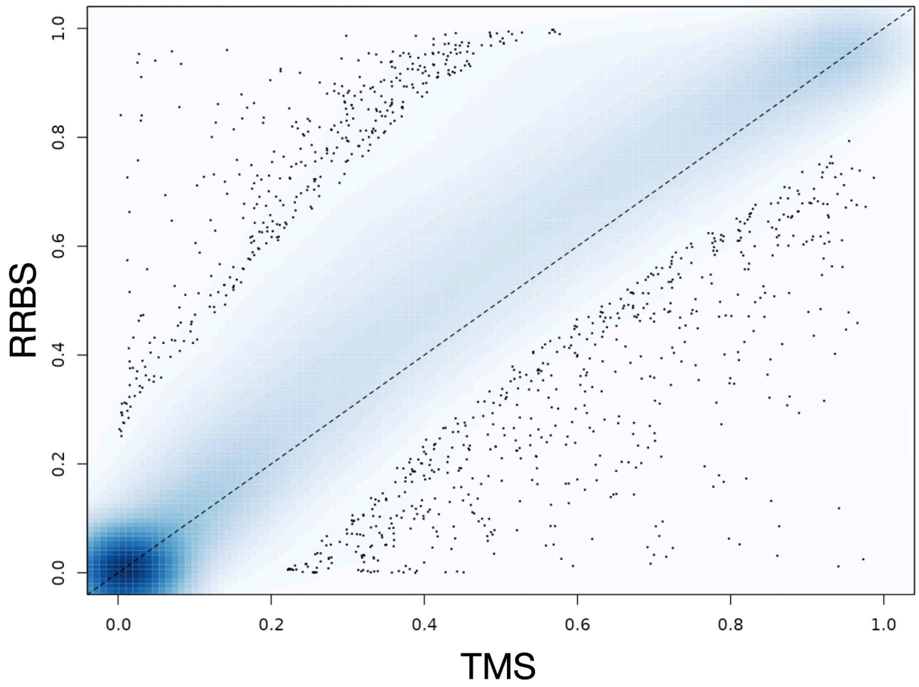


**Figure S10: Correlation in average site-level methylation between TMS and RRBS.** Site-level methylation averaged across 96 rhesus macaque samples processed using TMS and RRBS. Each point represents a site measured across both shearing methods and $R^2$ values were generated using linear modeling.

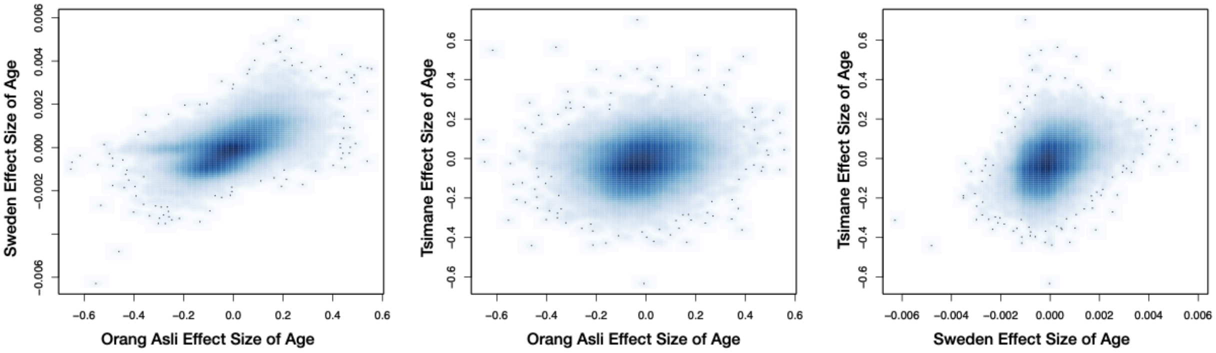


**Figure S11: Standardized effect sizes of age are correlated between populations living across very different environments.** Each plot shows the correlation of standardized effect sizes of age. Age effects were modeled at the CpG-level.

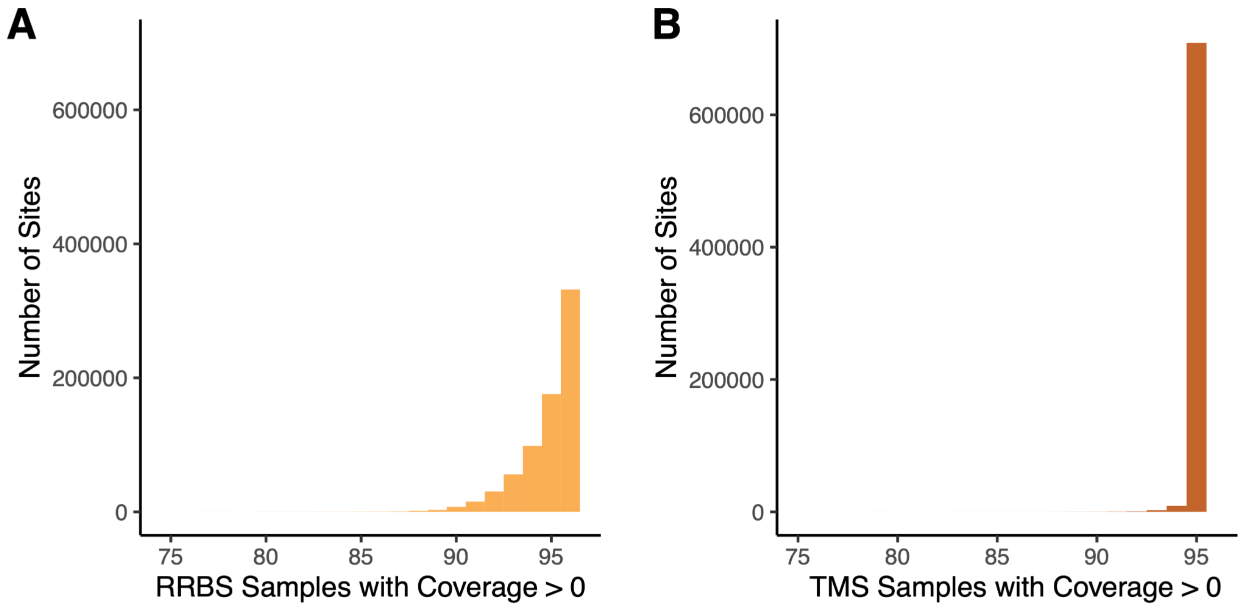


**Figure S12: Number of samples for which a site is covered across for datasets generated using (A) RRBS and (B) TMS.** Sites filtered for >5X coverage in >75% of samples processed using a given technology. A greater number of sites are covered consistently across all 96 samples using TMS compared to RRBS.

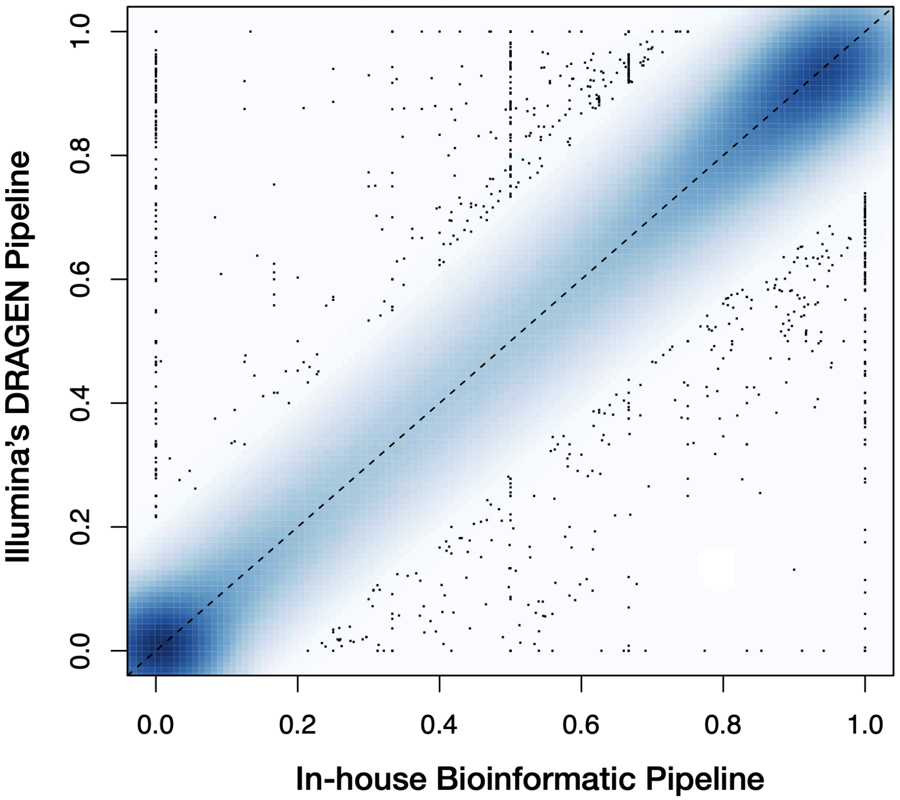


**Figure S13: Correlation in average site-level methylation between samples processed using Illumina's DRAGEN pipeline and our custom pipeline.** Each point represents the average methylation at a given site for 88 samples that were processed using both pipelines ($R^2 = 0.9972$).

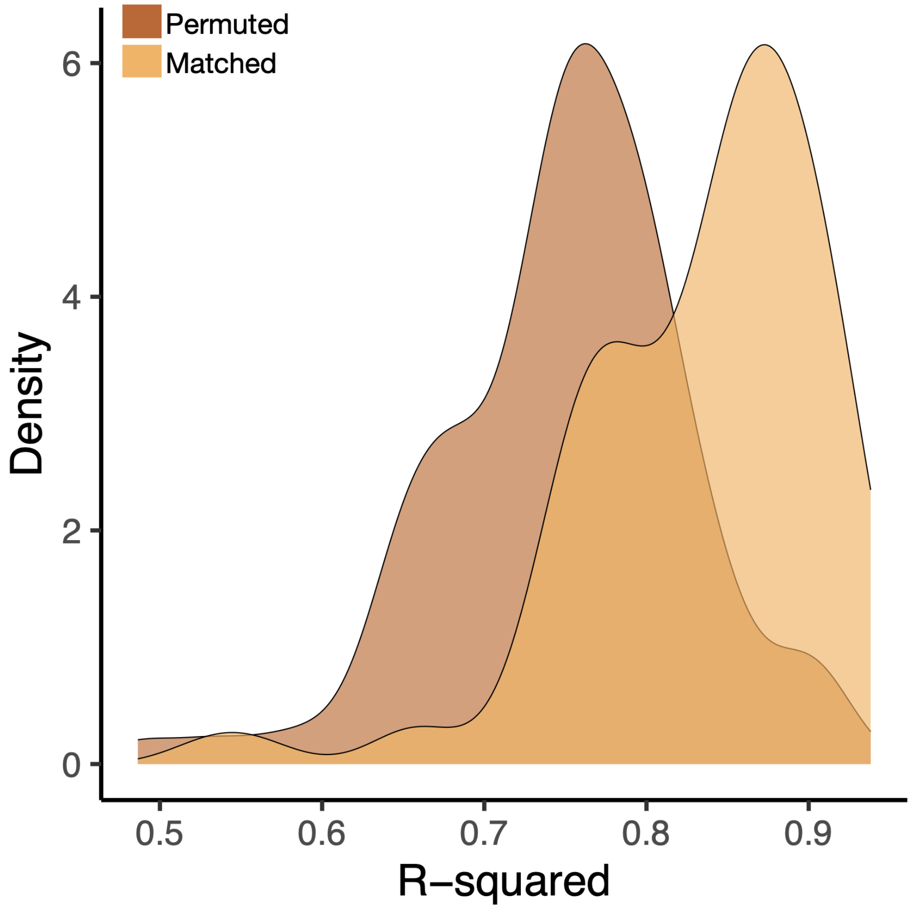


**Figure S14. Correlation in site-level methylation between TMS and RRBS data after the permutation of sample identifiers.** For samples processed using both TMS and RRBS, we assessed the correlation in site-level methylation for all sites after permuting sample ID randomly and compared them to non-permuted, or matched, sample IDs. $R^2$ values were generated using linear modeling. Using a t.test, we found a significant difference between the means of the two samples ($t = 7.6796$, p-value = $8.224 \times 10^{-13}$, mean of matched samples: 0.8345, mean of permuted samples: 0.7508).
